# Human tau promotes Warburg effect-like glycolytic metabolism under acute hyperglycemia conditions through modulating the homeostasis of protein-membrane association

**DOI:** 10.1101/2024.06.20.599836

**Authors:** Jinyi Yao, Zhenli Fu, Keying Li, Jingjing Zheng, Zicong Chen, Jiahao Xu, Guoqing Lai, Yaomin Huang, Jinsheng Huang, Guanying You, Shuangxue Han, Zhijun He, Qiong Liu, Nan Li

## Abstract

The neurofilaments formed by hyperphosphorylated tau is a hallmark of tau-related neurodegenerative disease, including Alzheimer’s disease, tau related FTDP-17, Pick’s disease, et al. However, the biological functions of tau and the physiological significance of its phosphorylation are still not fully understood. By using human tau (441 a.a.) transgenic (hTau) mice in which murine tau has been deleted simultaneously, murine tau knockout (Tau KO) mice and C57BL/6J (C57) mice, unexpectedly, we found that under acute hyperglycemia conditions, JNK but not previously reported GSK-3β mediated tau phosphorylation. Moreover, Akt, the upstream GSK-3β inhibitory kinase, was activated in a tau dependent manner. By comparing the membrane-associated proteome, we found that human tau influenced the homeostasis of protein-membrane association under acute hyperglycemia conditions. Of note, with respect to WT and Tau KO mice, the membrane-association of Krts, TFAM, TRAP1, mTOR et al, were strengthened by human tau. Whereas, the membrane-association of ribosomal proteins Rpls, proteasome proteins Psmds, and mitochondrial proteins, such as COXs, Ndufa1, Mtnt4, et al, were impeded by human tau. In vitro study showed that aerobic glycolysis was promoted in the presence of human tau, which maintained NAD^+^/NADH ratio. On the other hand, it restricted oxidative phosphorylation level, modulated the activity of SDH, and reduced ROS production upon challenging by high glucose. Furthermore, under acute high glucose conditions, the presence of human tau significantly augmented Akt activation, but inhibited 4EBP phosphorylation simultaneously, indicating that human tau is also involved in regulating the alternative activation of mTORC1/2. In summary, the current study revealed that human tau played an important role in regulating glycolytic metabolism under acute high hyperglycemia conditions, which is similar with the Warburg-effect, through influencing the homeostasis of protein-membrane association.

## Main

Tau was originally discovered as a microtubule binding protein. Recently, many studies also showed that tau played an important role in insulin signaling pathway. For example, tau deficient mice showed glucose intolerance and blunt response to insulin stimulation ^1–3^. The deficiency of tau also perturbed glucose-stimulated insulin secretion ^4^. Previous study demonstrated that tau inhibited the activity of phosphatase and tension homologue on chromosome 10 (PTEN). Therefore, loss of tau protein resulted in uncontrollable of 3,4,5-phosphatidyl inositol triphosphate (PIP3) dephosphorylation and the attenuation of insulin signaling ^5^. The observations from mice model demonstrated that murine tau closely relates with insulin signaling pathway. However, the biological function may strikingly different between murine tau and human tau, since the interactome of human tau showed that it also interacted with multitudinous mitochondrial proteins besides tubulin, including SGUCLG1, SUCLG2, SCL25A6, OXCT1, COX5B, VDAC2 and CYCS, et al. ^6^ Moreover, the overexpression of human tau resulted in mitochondrial elongation and the reduction of mitofusion 2 (MFN2) ubiquitination ^7^. In flies, the expression of human tau also affected the expression of Drp1 and Marf (the homolog of human MFN2)^8^. These studies illustrated that human tau is also involved in the biogenesis of mitochondria in neurocyte.

In the field of neurodegenerative disease research, many efforts had been made to demonstrate how hyperphosphorylation of tau was triggered by stress, such as amyloid-beta (Aβ), or hyperglycemia, and the subsequent neurotoxicity induced by the seeding, propagation of aggregated tau. However, little is known about the physiological meaning of tau phosphorylation itself. The biological function of human tau is quite vague, mainly because of its flexible structure and the obscure numerous interacting proteins. In the current study, we determined the role of tau phosphorylation in brain metabolisms by using acute hyperglycemia model, and monitor the influences of human tau on insulin signaling pathway and the functions of mitochondria through comparing human tau (441 a. a.) transgenic mice (hTau), which do not express murine tau, with C57BL/6J (C57), tau knockout (Tau KO) at the very the early stage of hyperglycemia conditions.

To our surprise, under the acute hyperglycemia conditions (<14 days), it was shown that the phosphorylation of tau at multiple sites were mediated by JNK, but not GSK-3β as known in the prolonged (>40 days) hyperglycemia mice model induced by STZ treatment^9^. On the other hand, the activation of Akt in brains, which is involved in mediating insulin signaling for glucose metabolism, was augmented by the presence of human tau under these conditions. Notably, the phosphorylation of Akt also accompanied with the decrease of pyruvate in brain of hTau mice. More importantly, the proteome from hippocampus showed that human tau maintained the homeostasis of membrane-associated proteins under the acute hyperglycemia conditions. In vitro study consistently showed that human tau strengthened high glucose induced Akt activation in a mTORC2 dependent manner, and facilitated aerobic glycolysis, thereby maintained NAD^+^/NADH ratio. Meanwhile, the oxidative phosphorylation level, as well as high glucose induced ROS production, were limited by the presence of human tau. These finding are also consistent with previous observations that neuronal somata exhibited increased aerobic glycolysis and reduced OXPHOS in both basal and activated state ^10^. Collectively, here we identified that human tau is involved in regulating protein-membrane association, especially those related with insulin signaling, oxidative phosphorylation, fatty acid oxidation, ribosome, and proteasome. By this way, it promoted a Warburg effect-like glycolytic metabolism under acute hyperglycemia conditions.

### Acute hyperglycemia induced tau phosphorylation in brain

Originally, the aim of this study was to monitor whether rapid tau phosphorylation induced by acute hyperglycemia would exhibit any neurotoxicity, especially through influencing mitochondria. Thus, human tau (441 a. a.) transgenic mice (hTau), tau knockout mice (Tau KO), and C57BL/6J mice (C57), which showed similar fasting blood glucose levels (**Fig. 1a**), were treated with intraperitoneal injection of STZ (150 mg/kg) to induce an acute hyperglycemia phenotype. This model have been widely used to study how hyperglycemia impacted on Aβ production, tau phosphorylation, as well as the recognizing ability ^9,11^. Previous reports demonstrated that hyperglycemia induced by STZ injection resulted in defects of insulin signaling, thus promoted tau phosphorylation in brain ^4,12^. However, to our surprise, at the very early stage of hyperglycemia, i.e. 3 days (**Fig. 1b**), and 7 days (**Fig. 1c**) after STZ injection, the level of fasting blood glucose of hTau mice were much lower than that of Tau KO mice and C57 mice, though it was significantly upregulated compared with non-STZ-injection group. To verify the effects of STZ treatment, we checked the levels of insulin (**Fig. 1d**), glucagon-like peptide-1 (GLP1) (**Fig. 1e**), and glucose dependent insulinotropic peptide (GIP) (**Fig. 1f**) in the serum. Interestingly, in non-fasting serum the levels of insulin, and GLP1 were not affected, but the level of GIP was significantly reduced by STZ treatment for 7 days. We further examined the phosphorylation levels of tau in the cortex 7 days after STZ injection (**Fig. 1g**). It was shown by the immunoblot that human tau was only detected in the cortex lysates from hTau mice by HT7 antibody (**Fig. 1h**). Tau-5 antibody, which recognize both human tau and murine tau, reflected the lack of tau in the cortex lysates from Tau KO mice (**Fig. 1i**). Interestingly, Tau-5 showed that STZ treatment reduced protein level of tau in the cortex, which did not reveal by HT7, indicating a discrepancy of these two antibodies. While, consistent with previous studies, the acute hyperglycemia induced increase of tau phosphorylation at multiple residues, including Thr181, Ser202/205, and Ser422 in the cortex of both C57 and hTau mice (**Fig. 1j-l**), and the phosphorylation of Thr231 in the cortex of hTau mice (**Fig. 1m**).

**Fig. 1.**
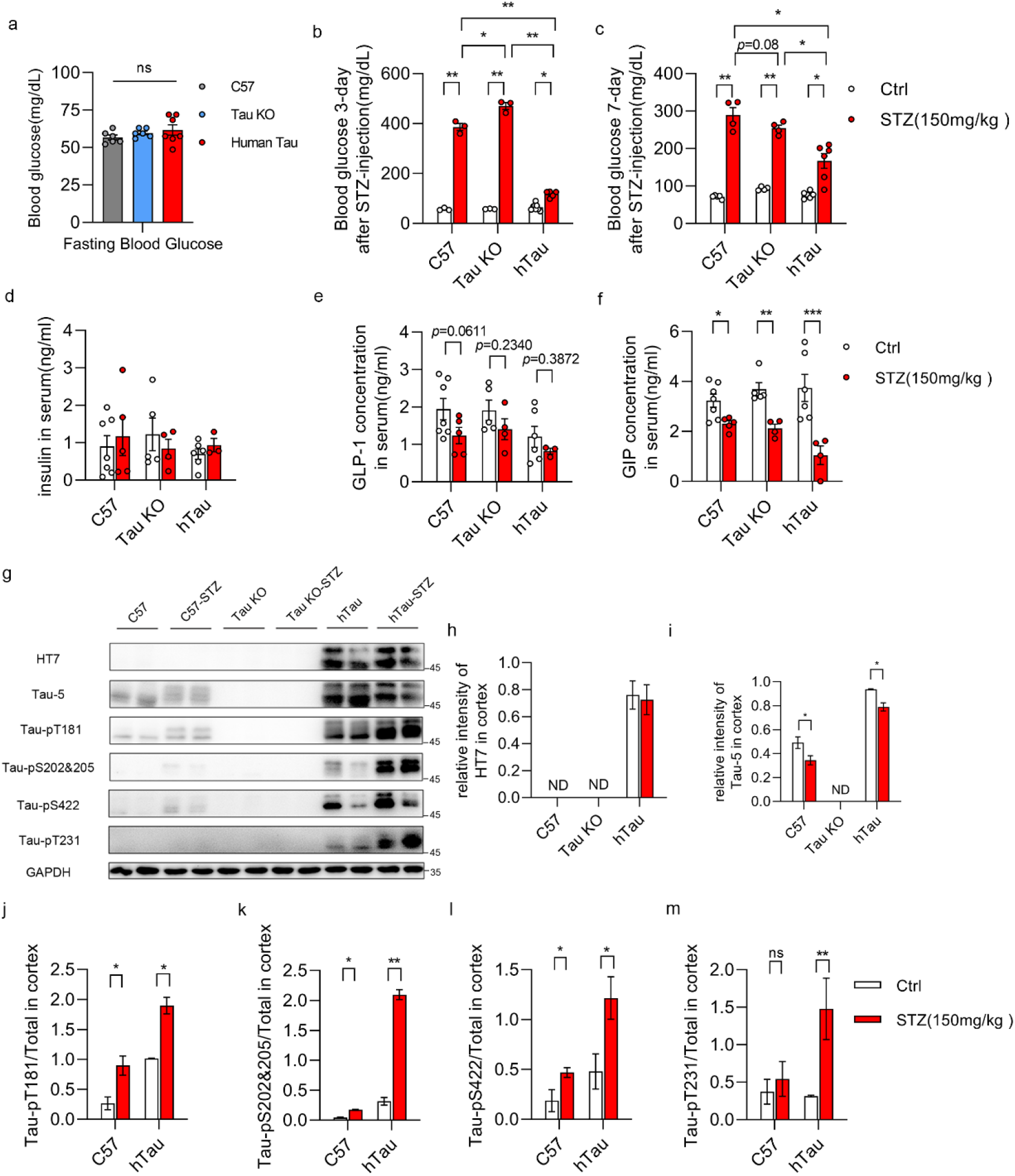
STZ injection induced acute hyperglycemia and tau phosphorylation. **a,** The levels of fasting-blood glucose in wild-type (WT, n=6), tau knockout (Tau KO, n=6), and human tau transgenic (hTau, n=7) mice. **b,** The levels of fasting-blood glucose in WT, Tau KO, and hTau mice 3 days after STZ (150 mg/kg) injection (WT, n=3. Tau KO, n=3. hTau, n=5) compared with control groups (WT, n=3. Tau KO, n=3. hTau, n=7). **c**, The levels of fasting-blood glucose in WT, Tau KO, and hTau mice 7 days after STZ (150 mg/kg) injection (WT, n=4. Tau KO, n=4. hTau, n=6) compared with control groups (WT, n=4. Tau KO, n=4. hTau, n=5). The levels of insulin (**d**), GLP-1 (**e**), and GIP (**f**) in non-fasting serum 7 days after STZ injection (WT, n=5. Tau KO, n=4. hTau, n=3) compared with control groups (WT, n=7. Tau KO, n=5. hTau, n=6). **g**, Representative immunoblot of the lysate of cortex 7 days after STZ injection (WT, n=4. Tau KO, n=4. hTau, n=4) compared with equal number controls. The level of human tau was detected by HT7 (**h**) (recognizes human tau specifically), he level of total tau was detected by tau-5 (**i**) (recognizes both human tau and murine tau), the levels of phosphorylated tau were detected by tau-pT181 (**j**), tau-pS204&205 (**k**), tau-pS422 (**l**), tau-pT231 (**m**), the protein level of GAPDH served as internal loading reference. The results are shown as the mean ±s.e.m., \**P*<0.05, \*\**P*<0.01, \*\*\**P*<0.001 by one-way ANOVA with Tukey’s test (**a**), two-way ANOVA with Tukey’s test (**b-f**) or two-tailed unpaired student’s *t*-test (**i-m**).

To determine whether the effects of STZ treatment on the levels of fasting blood glucose, and serum levels of insulin and GIP were repeatable, we employed 3xTg AD model mice, which also express 4R human tau ^13^ and their control wild-type (WT) mice. The level of fasting glucose of 3xTg AD mice were also significantly lower than that of WT mice 2 w after STZ injection (**Extended Data Fig. 1a**), though they were all significantly upregulated by STZ treatment compared with their non-treated controls, respectively. Interestingly, the levels of insulin (**Extended Data Fig. 1b**) and GIP (**Extended Data Fig. 1c**) in non-fating serum of 3xTg AD mice were significantly lower than that of WT mice. In addition, 2 w after STZ injection the level of insulin in the serum of WT mice was decreased compared with non-treated WT mice (**Extended Data Fig. 1b**), which was not seen in 7 days STZ treatment C57 mice, suggesting that the downregulation of insulin by STZ treatment is behind of GIP decrease. While, the level of GLP1 was decreased in both WT and 3x Tg AD mice compared with their control groups, respectively (**Extended Data Fig. 1c**). We further examined the effect of hyperglycemia on tau phosphorylation in the cortex of WT mice and 3x Tg AD mice as well (**Extended Data Fig. 1d**), and found that STZ treatment reduced the protein level of tau in the cortex of WT mice (**Extended Data Fig. 1e**) but not in 3x Tg AD mice. We assume that may because of the overexpression of human tau in these mice (**Extended Data Fig. 1f**). Though WT mice did not express human tau (specifically recognized by tau-13 antibody), the murine tau could be recognized by the antibodies for tau phosphorylation as well. Consistently, the hyperglycemia induced by STZ treatment for 2 w also resulted in increased tau phosphorylation at Thr181, Ser422, Ser404, and Ser396 in the cortex from 3x Tg AD mice (**Extended Data Fig.1g-j**), ant tau phosphorylation at Thr181 and Ser296 in the cortex of WT mice (**Extended Data Fig.1g, j**).

### JNK inhibitor attenuated tau phosphorylation induced by high glucose

Previous studies demonstrated that STZ induced long-term (>40 days) hyperglycemia caused tau phosphorylation through activating GSK-3β, due to the deficiency of insulin and subsequent inactivation of Akt in brain ^4,12^. To determine whether GSK-3β is also responsible for tau phosphorylation at the acute hyperglycemia conditions. We monitored the phosphorylation levels of Akt and GSK-3β by immunoblot (**Extended Data Fig. 2a**). Unexpectedly, the phosphorylation of Akt at Ser473 was significantly increased (**Extended Data Fig. 2a, b**), and the inhibitory phosphorylation of GSK-3β at Ser9 was also enhanced in the cortex of hTau mice 7 days after STZ injection (**Extended Data Fig. 2a, c**). These results, however, indicated that GSK-3β may not account for tau phosphorylation under the acute hyperglycemia conditions. Then, we further examined the phosphorylation levels of CDK5, PKA, CAMKII, AMPK, and JNK, as well as the protein levels of CDK5 activator, P35/P25, in the cortex by immunoblot. Interestingly, it was shown that the phosphorylation level of CDK5 (**Extended Data Fig. 2a, d**) and the protein level of p35 (**Extended Data Fig. 2a, e**) were not changed in the cortex 7 days after STZ injection. Moreover, the phosphorylation levels of CAMKIIα/β were inhibited in the cortex from Tau KO and hTau mice (**Extended Data Fig. 2a, f, g**). Strikingly, the phosphorylation level of PKA was significantly higher in the cortex from Tau KO mice compared with that of C57 and htau mice. 7 days after STZ injection, it was upregulated in C57 mice, but decreased in Tau KO mice compared with non-treated groups (**Extended Data Fig. 2a, h**), respectively. While, the phosphorylation level of AMPK was significantly reduced in the cortex from hTau mice 7 day after STZ treatment compared with non-treated mice (**Extended Data Fig. 2a, i**). Among these kinases, only JNK was activated in the cortex of STZ treated hTau mice compared with non-treated hTau mice, indicating JNK may be contribute to triggering tau phosphorylation under acute hyperglycemia conditions.

However, these results are conflict with the previous studies ^4,12^. We presumed that this may because of the different time course between our experiment and others’. Therefore, we also compared the phosphorylation levels of Akt and GSK-3β in the cortex of WT mice and 3xTg AD mice 2 w and 6 w after STZ injection with non-treated groups, respectively. As expected, the phosphorylation levels of Akt at Ser473, and GSK-3β at Ser9 were decreased in the cortex from WT mice (**Extended Data Fig. 3a-c**) and 3xTg AD (**Extended Data Fig. 3g-i**) mice after long-term (6 w) STZ treatment, respectively. More importantly, we also found that the protein level of insulin degrading enzyme (IDE) was decreased upon STZ treatment in the cortex of either WT (**Extended Data Fig. 3a, e**) mice or 3x Tg AD mice (**Extended Data Fig. 3g, k**), and the phosphorylation of mTOR was increased in the cortex of 3x Tg AD mice (**Extended Data Fig. 3j**) though was not in WT mice (**Extended Data Fig. 3d**). Astonishingly, the protein level of insulin was significantly increased in both of the long-term hyperglycemia groups (**Extended Data Fig. 3a, f, g, l**), implying that the inactivation of Akt is caused by insulin resistant instead of insulin deficiency. Additionally, though the phosphorylation levels of JNK and p38 in the cortex of WT mice were not severely affected by STZ treatment (**Extended Data Fig. 4a-c**), we speculated that the transient activation of JNK by hyperglycemia may sufficient for triggering tau phosphorylation at certain residues as seen in **Fig. 1g** and **Extended Data Fig. 1d**. However, in 3x Tg AD mice, 2 w after STZ injection, phosphorylation levels of JNK and p38 were increased, but 6 w after STZ injection they were significantly reduced (**Extended Data Fig. 4f-h**) in the cortex compared with that from non-treated mice. Interestingly, the phosphorylation levels of CaMKII α/β were further reduced in both of the long-term STZ treatment groups (**Extended Data Fig. a, d, f, j**), indicating that long-term hyperglycemia triggered impairment of neuroplasticity ^14^, which was not dependent on the existence of human tau.

To confirm whether JNK was responsible for tau phosphorylation under acute hyperglycemia conditions in vitro, we used a human tau (441 a. a.) overexpressing HEK293 cell line (239tau), and found that high glucose (60 mM) treatment for 24 h significantly induced JNK, p38, and Erk1/2 phosphorylation in these cells (**Fig. 2a-d**), but only triggered Erk1/2 phosphorylation in normal HE293 cells (293) (**Fig. 2a, d**). More importantly, high glucose also resulted in tau phosphorylation (**Fig. 2e, f-i**), as well as tau dependent Akt, GSK-3β, and mTOR phosphorylation (**Fig. 2e, k-m**). Whereas, when JNK inhibitor SP600125 (10 mM) were added together with high glucose, the phosphorylation of tau was blocked efficiently along with the inactivation of JNK (**Fig. 2e, f-i**). Meanwhile, JNK inhibitor also significantly arrested high glucose induced Akt, GSK-3β, and mTOR phosphorylation in 293tau cells (**Fig. 2e, j-m**). These results indicated that JNK mediated tau phosphorylation under high glucose treatment, and the phosphorylation of tau is also essential for triggering mTOR and Akt activation under the stress. However, it is well known that mTOR participates in forming 2 complexes. Together with Raptor, mLST8, et al., they constitute mTORC1 which generates negative feedback to insulin signaling subsequently results in inactivation of Akt, but induces phosphorylation of 4EBP-1, S6K. Together with Rictor, mLST8, and mSin1, et al., they form mTORC2 which facilitates phosphorylation of Akt on Ser473 residue ^15^. To demonstrate which complex is responsible for the activation of Akt in tau expressing HEK293 cells, we compared the phosphorylation levels of 4EBP-1 and Akt in tau expressing HEK293 cells and normal HEK293 cells that treated with high glucose for 0-24 h. The immunoblot results showed that high glucose induced 4E-BP-1 phosphorylation only in normal HEK293 cell, whereas, it led to Akt activation in both cell lines. However, in normal HEK293 cells, the phosphorylation of Akt was diminished soon, whereas it was prolonged by the presence of tau (**Extended Data Fig. 5a**). When INK-128, Rapamycin, JR-AB2-011, the inhibitor of mTORC1/2, mTORC1, and mTORC2, respectively, were added together with high glucose. It was shown that INK-128 and JR-1B2-011 efficiently inhibited high glucose induced Akt phosphorylation in tau expressing HEK293 cells. Meanwhile, INK-128 and Rapamycin significantly reduced high glucose triggered phosphorylation of 4E-BP-1 in normal HEK293 cells (**Extended Data Fig. 5b**). These results suggested that human tau promoted the alternative activation of mTORC2 under high glucose conditions, thereby prolonged high glucose induced Akt activation. This is similar to the alterative activation of mTORC2 by posttranslational modification of Rictor ^16,17^, which is also a tau interactor. However, the detail of underlying mechanisms needs to by further studied.

**Fig. 2.**
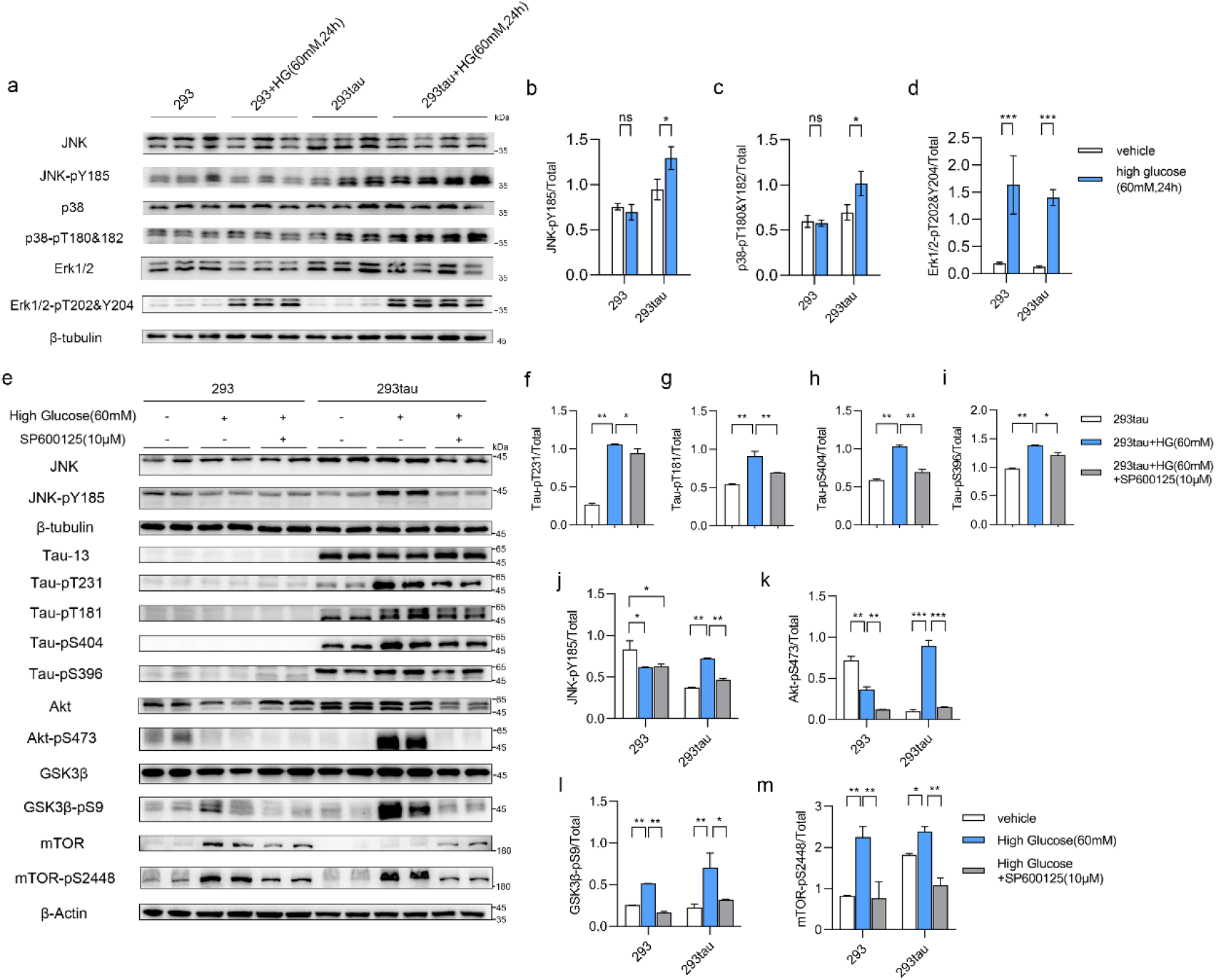
Inhibition of JNK arrested high glucose induced tau phosphorylation and tau dependent Akt and mTOR activation. **a,** Immunoblot analysis of MAPKs activation in normal HEK293 cells (293) and tau expressing HEK293 cells (293tau) that were cultured in high glucose (60 mM) medium for 24 h compared with that cultured in normal medium. The results are shown as the mean±s.e.m. of duplicates for JNK-Py185/JNK (**b**), p38-pT180&182/p38 (**c**), and Erk1/2-pT202&204/Erk (**d**). **e,** Representative immunoblot analysis the phosphorylation levels of JNK, tau, Akt, GSK3β, and mTOR and in normal HEK293 cells (293) and tau expressing HEK293 cells (293tau) after 24 h of high glucose treatment (60 mM),combined with adding JNK inhibitor SP600125 (10 μM) in high glucose medium compared with that cultured in normal medium. The results are shown as the mean±s.e.m. of 3 independent experiments for tau-pT231/tau (**f**), tau-pT181/tau (**g**), tau-pS404/tau (**h**), tau-pS396/tau (**i**), JNK-pY185/JNK (**j**), Akt-pS473/Akt (**k**), GSK3β-pS9/GSK3β (**l**), and mTOR-pS2448/mTOR (**m**). \**P*<0.05, \*\**P*<0.01, \*\*\**P*<0.001 by two-tailed unpaired student’s *t*-test (**b-d**) or one-way ANOVA with Tukey’s test (**f-m**).

### Human tau promoted aerobic glycolysis under acute high glucose conditions

Under the acute hyperglycemia conditions induced by STZ injection, human tau enhanced Akt and mTOR activation as seen above. Due to the important roles of mTOR and Akt in mediating insulin signaling and facilitating glycolysis and tricarboxylic acid (TCA) cycle ^18^, we were curious whether tau exerts any influence on glycolysis under these conditions. Therefore, we examined the levels of pyruvate and lactate, the main products of glycolysis, in the cortex of mice brains. The level of pyruvate was significantly increased in the cortex from Tau KO mice upon STZ injection for 7 days. However, it was significantly lower in the cortex from hTau mice under the acute hyperglycemia conditions compared with those from C57 and Tau KO mice with the same treatment (**Fig. 3a**). The pyruvate produced by glycolysis will be further subjected to dehydrogenation to form lactate and replenish NAD^+^, or to form acetyl-CoA to participate in TCA cycle or biosynthesis, through the catalyzation of LDHA or PDK, respectively. Thus, we further compared the levels of lactate in the cortex. It was shown that in all three genotype mice it was upregulated to a similar level 7 days after STZ injection (**Fig. 3b**).

**Fig. 3.**
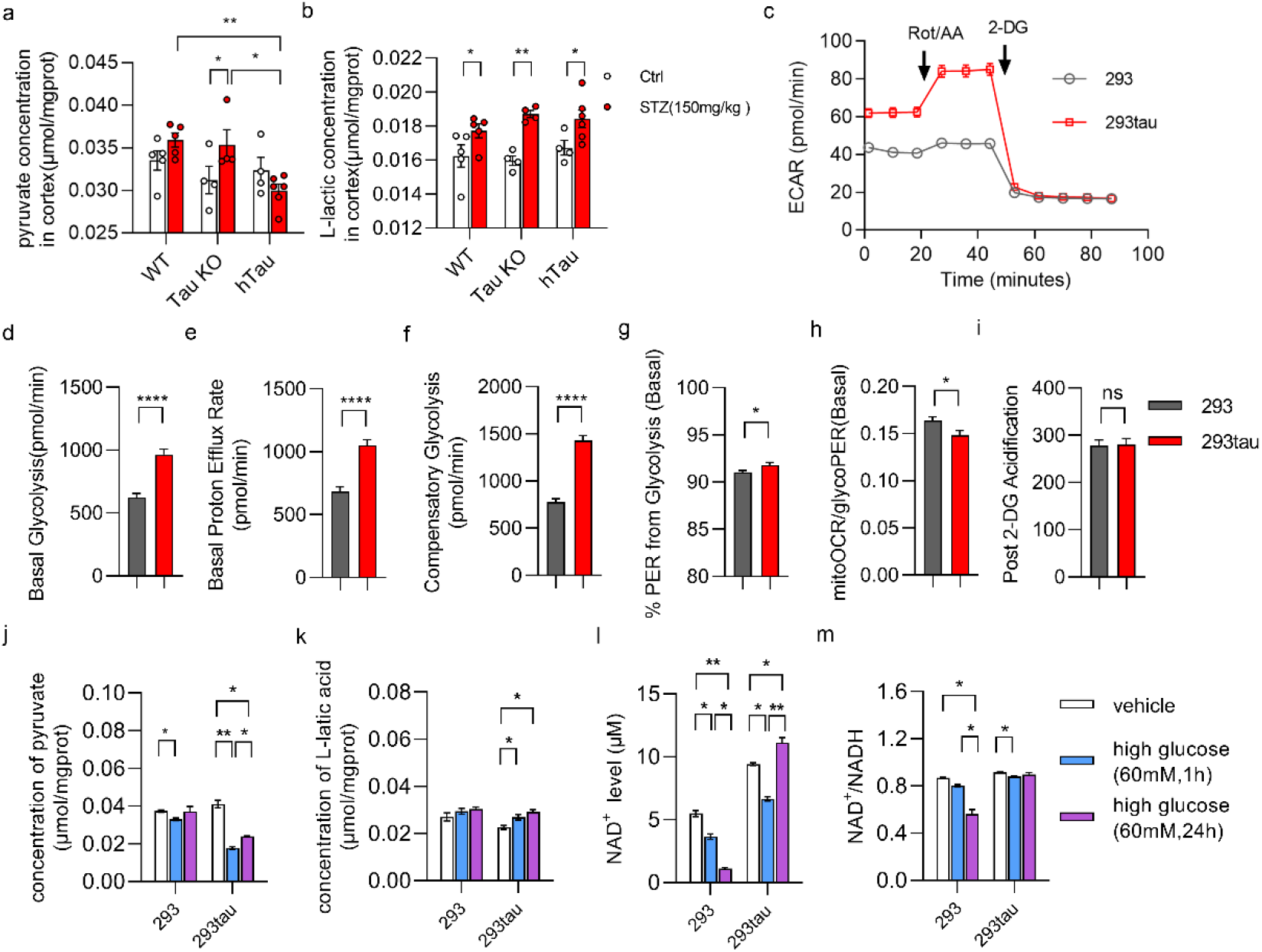
Human tau promoted aerobic glycolysis The concentration of pyruvate. (**a**) and lactate (**b**) in the lysates of cortex 7 days after STZ injection (WT, n=5. Tau KO, n=4. hTau, n=6) compared with control groups (WT, n=5. Tau KO, n=5. hTau, n=6). **c**, Representative bar graph of Extracellular acidic rate (ECAR) trace of normal HEK293 cells (293) and tau expressing HEK293 cells (293tau) measured by using the Seahorse apparatus. The levels of basal glycolysis (**d**), basal proton efflux rate (PER) (**e**), compensatory glycolysis (**f**), %PER from glycolysis (**g**), mitoOCR/glycoPER (**h**), and extracellular acidification rate after the hexokinase inhibitor 2-DG have been added (**i**) were compared between the two cell lines. The concentration of pyruvate (**j**) and lactate (**k**) in the lysates of normal HEK293 cells and tau expressing HEK293 cells that were treated with high glucose (60 mM) for 1 h or 24 h compared with that from normal medium cultured cells, respectively. The level of NAD^+^ (**l**) and the ratio of NAD^+^/NADH (**m**) in the lysates of normal HEK293 cells and tau expressing HEK293 cells that were treated with high glucose (60 mM) for 1 h and 24 h compared with that from normal medium cultured cells, respectively. The results are shown as the mean± s.e.m., \**P*<0.05, \*\**P*<0.01, by two-way ANOVA with Tukey’s test (**a, b**), two-tailed unpaired student’s *t*-test (**d-i**), one-way ANOVA with Tukey’s test (**j-m**). The experiments were repeated for 3 times by using different batch of cells.

It seems that the acidic circumstance could be restrained within a certain extend in the central nervous system. Thus, we decided to detect the impact of tau on the capability of glycolysis by performing extracellular acidification rate (ECAR) analysis in vitro. It was clearly shown that the basal glycolysis in human tau expressing HEK293 cells (293tau) was significantly higher than that in normal HEK293 (293) (**Fig. 3c, d**) as also reflected by the proton efflux rate (PER) (**Fig. 3e**). The level of compensatory glycolysis in tau expressing HEK293 cells was also significantly greater than that in HEK293 cells, when oxidative phosphorylation was inhibited by rotenone/antimycin A (Rot/AA) (**Fig. 3f**). The %PER from glycolysis was also higher in tau expressing HEK293 cells (**Fig. 3g**). However, the mitochondrial oxygen consumption rate (mitoOCR)/ glycolysis PER was less in these cells compared with normal HEK293 cells, indicating a less oxygen consumption for glucose metabolism in tau expressing cells (**Fig. 3h**). When glycolysis was inhibited by adding hexokinase inhibitor 2-DG, the ECAR were stopped in both cell lines (**Fig. 3i**), illustrating that the extracellular protons were indeed coming from aerobic glycolysis.

As mentioned above, pyruvate is subjected to dehydrogenation to form lactate in the process of aerobic glycolysis, which replenished NAD^+^ simultaneously. The in vivo data and ECAR results demonstrated that tau promoted Akt activation and aerobic glycolysis. Thus, we further detected the levels of pyruvate, lactate, and NAD^+^ in cells under high glucose (60 mM) stress. It was shown that the level of pyruvate in tau expressing HEK293 cells was significantly reduced 1 h and 24 h after high glucose had been added comparing with those were not treated with high glucose (**Fig. 3j**). Meanwhile, the level of lactate was increased along with the reduction of pyruvate (**Fig. 3k**). More importantly, 1 h after high glucose treatment, the level of NAD^+^ was decreased in both tau expressing HEK293 and normal HEK293 cells compared with control group, respectively. However, 24 h after high glucose treatment, the level of NAD^+^ was increased significantly in tau expressing HEK293 cells, in contrast, it went down continuously in normal HEK293 cells (**Fig. 3l**). The ratio of NAD^+^/NADH was also significantly reduced in normal HEK293 cells upon high glucose treatment for 24 h, but it was maintained by the present of tau (**Fig. 3m**). These results demonstrated that human tau promoted aerobic glycolysis under high glucose conditions, which further transform pyruvate into lactate, so that the level of NAD^+^ was replenished efficiently.

### Human tau took part in regulating the homeostasis of membrane-associated protein

We presumed that tau mediated activation of Akt attribute to the alternatively activation of mTORC2. It had been illustrated that the activation of mTOR is dependent on the association of mTOR to the plasm membrane ^19^. In addition, tau was also found to interact with lipid membranes ^20^. Thus, for seeking the mechanism underlying tau regulating kinase activity such as Akt, mTOR, we decided to compare the membrane-associated proteome from the hippocampus of different genotype mice. The membrane associated protein were isolated from hippocampus (**Extended Data Fig. 6a**) and subjected to TMT-iTRQ-MS. The proteome of membrane-associated proteins showed that when comparing with C57 mice, in the hippocampus of Tau KO mice 90 proteins were significantly more abundant in the membrane extract, including Zfyve19, Rpls, Psmds, et al., however, 90 proteins were significantly less associated with membrane, including Nnt, Mapt (tau), Trap1, GSK3β,et al. (**Fig. 4a**). When comparing with C57 mice, 24 proteins were more abundant in the membrane extract, including Zfyve19, Mapt, TFAM, and Krt13, et al., however, 39 proteins were less associated membrane, including Nnt, Ndufb1, Psmd5 et al., in the hippocampus of hTau mice (**Fig. 4b**). These results indicated that Nnt, Zfyve19 may not be affected by tau protein. Instead, we suspected that the human tau transgenic mice and tau knockout mice may derived from a strain of B6 mice, which do not express Nnt gene ^21^. When compared with Tau KO mice, on the other hand, the proteome showed that 209 proteins, including Mapt, RNF19a, Krts, Ercc1, TFAM, TRAP1, mTOR, et al. were more abundant on the membrane in the hippocampus of hTau mice. Interestingly, the membrane association of the JNK activator Map2k4 was also increased by human tau (**supplementary material 1**), which may explain acute hyperglycemia triggered JNK phosphorylation in brains of tau expressing mice. Whereas, 179 proteins, including Rpls, Psmds, Ndufb1, Ndufaf3, Cox5a1, Cox7a2, HSPs, et al. were less enriched on membrane in hippocampus of hTau mice (**Fig. 4c**). KEGG analysis demonstrated that deficiency of tau resulted in the elevation of ribosome related protein but the decline of TCA cycle related protein in the membrane extract of hippocampus compared with C57 mice (**Fig. 4d**). It also showed that the proteins related to oxidative phosphorylation were less abundant on the membrane in hippocampus from the hTau mice compared with both c57 and Tau KO mice (**Fig. 4e, f**), indicating that human tau may interfere with oxidative phosphorylation. Protein-protein interaction (PPI) network analysis illustrated that trap1, which is also known as a mitochondrial chaperone and involved in regulating respiratory capacity ^22^, was the top hub node in the differential membrane-associated proteome of hippocampus between Tau KO and C57 mice (**Fig. 4g**). While, TFAM and mTOR played central roles in the differential membrane-associated proteome of hippocampus between hTau and C57 mice, and between hTau and Tau KO mice, respectively.

**Fig. 4.**
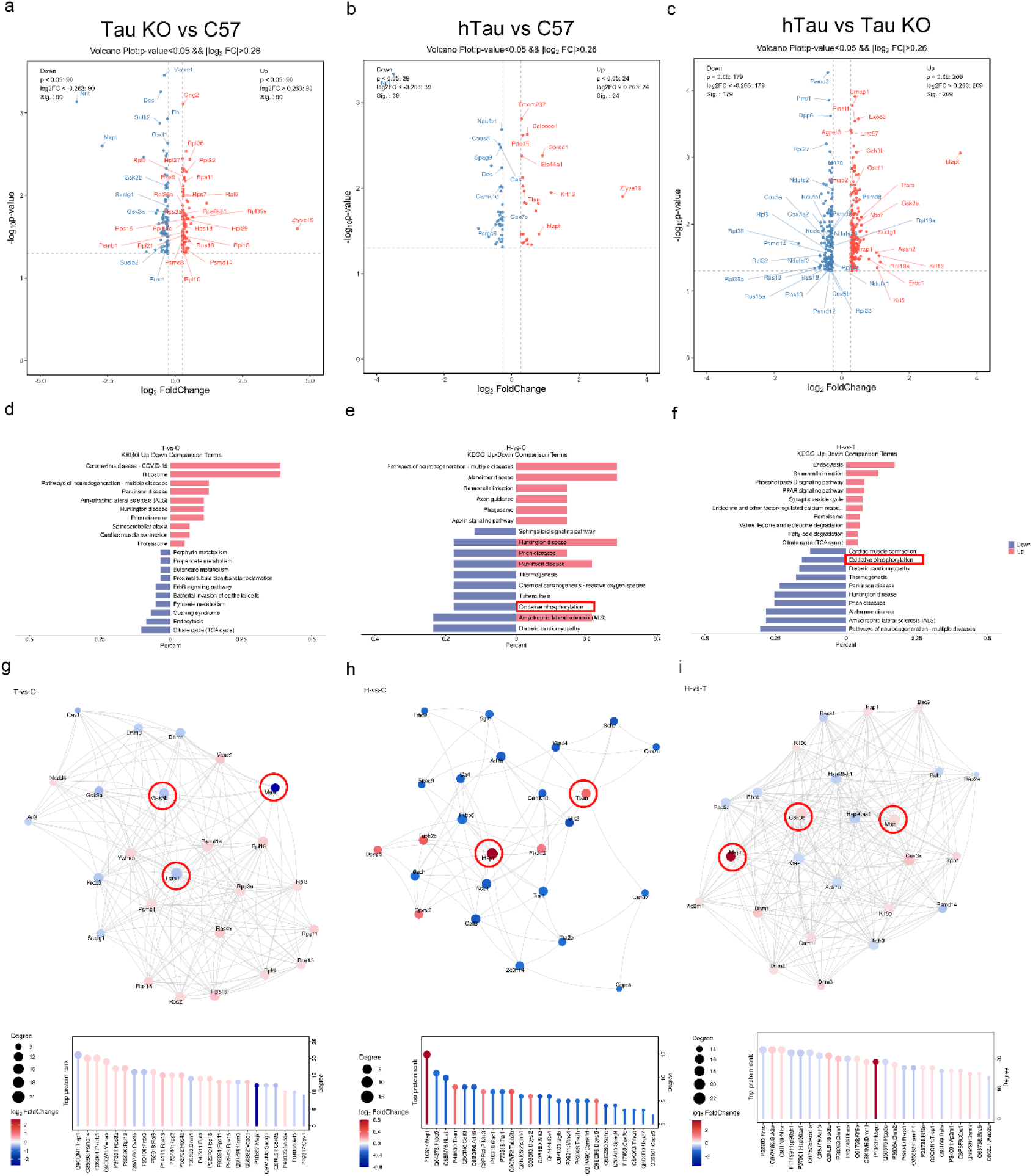
Tau altered the state of membrane-associated proteins in the hippocampus. Volcano plots of the membrane-associated proteins (*P*<0.05, log_2_ Fold Change>0.26) in the hippocampus from 4 months old male mice (**a-c**). **a**, By comparing with C57 mice (n=4), 90 proteins were significantly more abundant and 90 proteins were significantly less abundant in association with membrane in the hippocampus from tau KO mice (n=4). **b**, By comparing with C57 mice (n=4), 24 proteins were significantly more enriched and 39 proteins were significantly less enriched in association with membrane in the hippocampus from hTau mice (n=4). **c**, By comparing with Tau KO mice (n=4), 209 proteins were significantly more plentiful and 179 proteins were significantly less plentiful in association with membrane hippocampus from hTau mice (n=4). KEGG analysis of the significantly regulated membrane-associated protein (**d-f**). **d**, Compared with C57 mice, murine tau knockout upregulated ribosome, neurodegenerative diseases related proteins, but downregulated TCA cycle, pyruvate metabolism related proteins in associated with membrane. **e**, Compared with C57 mice, human tau upregulated axon guidance, phagosome related proteins, but downregulated oxidative phosphorylation, thermogenesis related proteins in associated with membrane. **f**, Compared with murine tau knockout mice, human tau upregulated endocytosis, peroxisome, and fatty acid degradation related proteins, but downregulated oxidative phosphorylation, thermogenesis protein in association with membrane. Protein-protein interaction (PPI) network retrieved from the STING databases predicted the hub protein (**g-i**). **g**, In the differential membrane-associated protein network between tau KO and C57 mice, TRAP1, GSK3β, et al. played central roles. **h**, In the differential membrane-associated protein network between human tau and C57 mice, MAPT, TFAM, et al. were the top rank hub. **i**, In the differential membrane-associated protein network between hTau and tau KO mice, Kras, GSK3β, and mTOR were predicted as the most important hub proteins.

We further studied the discrepancy of membrane-associated protein in hippocampus under acute hyperglycemia conditions among the above three genotype mice. Strikingly, in the hippocampus of hTau mice, 7 days after STZ injection, only 2 proteins were upregulated and 8 proteins were downregulated significantly in association with membrane compared with non-treated hTau mice (**Extended Data Fig. 6b**). However, under the same conditions, in the hippocampus of C57 mice, there were 101 and 84 membrane associated proteins which were upregulated and downregulated comparing with non-treated C57, respectively (**Extended Data Fig. 6c**). In addition, acute hyperglycemia resulted in 196 proteins more abundant on the membrane, but led to 181 proteins less combining with membrane in hippocampus of Tau KO mice (**Extended Data Fig. 6d**). KEGG analysis indicated that the membrane-associated proteins that were elevated by acute hyperglycemia in C57 and Tau KO mice are related with neurogenerative disease, such as Alzheimer’s disease, Parkinson disease, Huntington disease (**Extended Data Fig. 6e, f**). This is consistent with the observations that these diseases are accompanied with dysregulation of glycose metabolism ^23^. Remarkably, the membrane association of oxidative phosphorylation related proteins were also augmented by acute hyperglycemia in the hippocampus of C57 and Tau KO mice, but not in hTau mice (**Extended Data Fig. 6e, f**). PPI network analysis showed that the downregulation of Lrpprc, and mTOR,which are all involved in regulating glycolysis ^24^, played important role in the differential membrane-associated proteome under acute hyperglycemia conditions, in WT and tauKO mice, respectively (**Extended Data Fig. 6g, h**). Collectively, these results indicated that human tau possesses the ability of maintaining the homeostasis of protein-membrane association under acute hyperglycemia conditions, especially for the proteins that are involved in oxidative phosphorylation. It directly or indirectly cooperated with the proteins that are involved in regulating glycolysis and mitochondria biogenesis, including TFAM, TRAP1, mTOR, GSK3β, et al..

### Human tau restricted oxidative phosphorylation and reduced ROS production under acute high glucose conditions

As the membrane-associated proteome shown, many proteins that are involved in oxidative phosphorylation were less abundant in the membrane extract of hippocampus from hTau mice compared with that from Tau KO mice, including Mtnds, Ndufs, QCRs, Coxs, (**Fig. 5a**). These results indicated that human tau (441 a.a.) may impede the functions of electron transport chain on the membrane of mitochondria. Therefore, we performed OCR (**Fig.5 b**) to compare the level of oxidative phosphorylation in tau expressing HEK293 (293tau) and normal HEK293 cell (293). As expected, the levels of basal respiration (**Fig. 5c**), maximal respiration (**Fig. 5d**), as well as the ATP production of tau expressing HEK293 cells were significantly lower than that of normal HEK293 cells as indicated by the less of O_2_ consumption. The proton leak, which switches the potential of proton to thermogenesis without ROS production, was restricted by tau as well (**Fig. 5e**). This is also consistent with the increased coupling efficiency in tau expressing HEK293 cells compared with normal HEK293 cells (**Fig.5 g**). However, it was also shown that the non-mitochondrial oxygen consumption (**Fig.5 h**) was increased in tau expressing HEK293 cells compared with normal HEK293 cells. We speculated that this is due to the upregulation of fatty acid oxidation as indicated by the membrane-association proteome which showed many proteins related to peroxisome (**Extended Data Fig. 7a**) and fatty acid oxidation were more abundant in membrane extract in the hippocampus from hTau mice compared with that from Tau KO mice (**Extended Data Fig. 7b**).

**Fig. 5.**
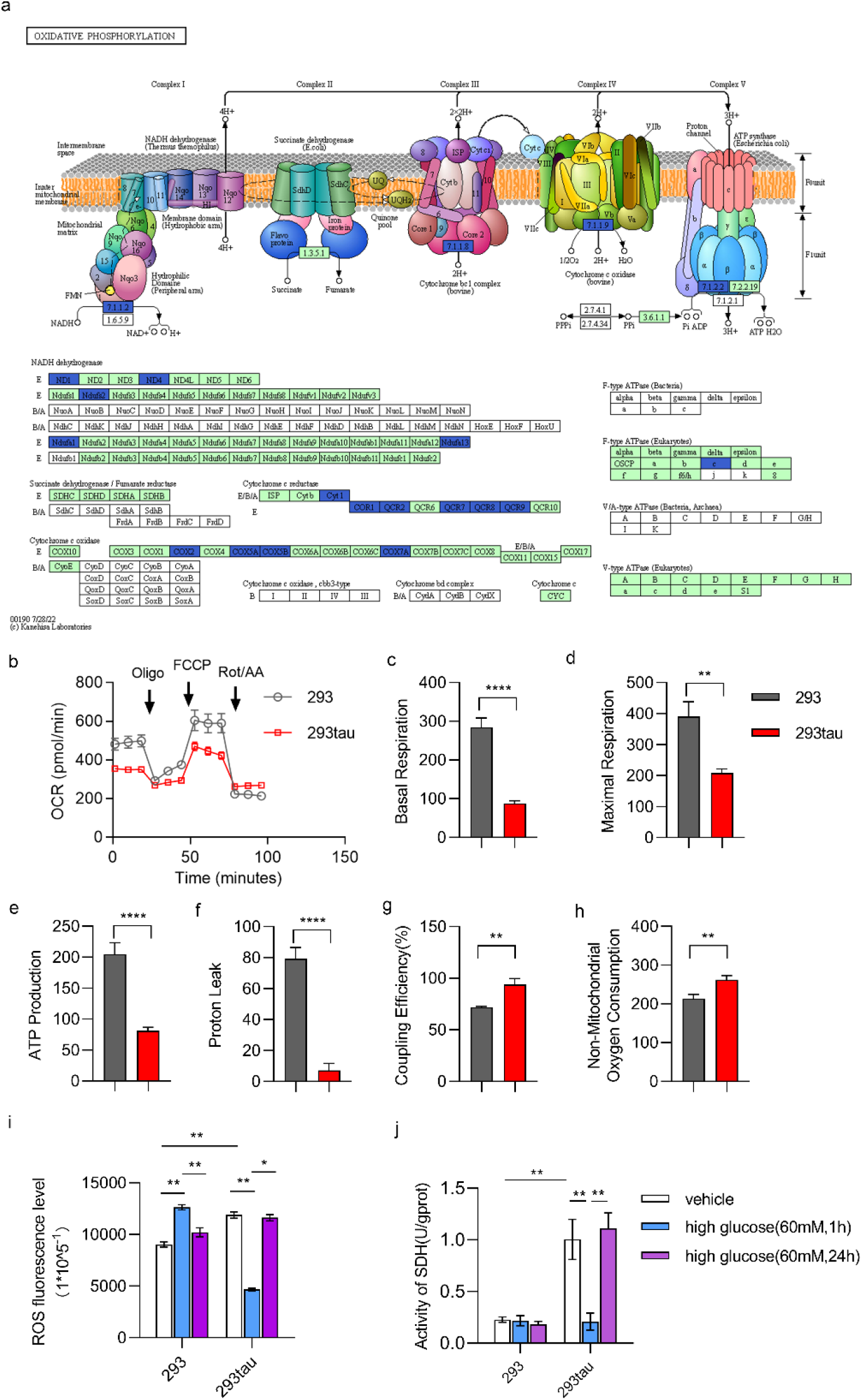
Human tau restricted oxidative phosphorylation and inhibited acute high glucose induced ROS production. **a**, The membrane associated proteome showed that the components of NADH dehydrogenase, cytochrome c reductase, cytochrome c oxidase, and F-type ATPase were less abundant in association with membrane in hippocampus from hTau mice comparing with that from tau KO mice. **b**, Representative graph of OCR from normal HEK293 cells (293) and tau expressing HEK293 cells (293tau) measured by using the Seahorse apparatus. The oxygen consumption for basal respiration (**c**), maximal respiration (**d**), ATP production (**e**), non-mitochondrial (**f**), proton leak (**g**), as well as the coupling efficiency (**h**) were calculated and shown as the mean±s.e.m.. **i**, The level of ROS in normal HEK293 and tau expressing HEK293 cells that were treated with high glucose (60 mM) for 1 h and 24 h were measured and compared with normal medium cultured cells, respectively. The fluorescence was normalized to 10^-5^ cells and shown as the mean ± s.e.m.. **j**, The activity of SDH in the lysates of normal HEK293 and tau expressing HEK293 cells that were treated with high glucose (60 mM) for 1 h or 24 h were measured and compared with normal medium cultured cells, respectively. The results were normalized to per gram of protein and shown as the mean±s.e.m., \**P*<0.05, \*\**P*<0.01, \*\*\**P*<0.001 by two-tailed unpaired student’s *t*-test (**c-h**) or two-way ANOVA with Tukey’s test (**i, j**). The experiments were repeated for 3 times by using different batch of cells.

Since the oxidative phosphorylation is also a main source of ROS. We further examined the level of ROS under acute high glucose conditions. To our surprise, the results showed that the level of ROS in tau expressing HEK293 cells was higher than that in normal HEK293 cells. We assumed that this may also attribute to the augment of fatty acid oxidation in tau expressing HEK293 cells. However, as expected, when challenged with high glucose for 1 h, the level of ROS was significantly increased in HEK293 cells, in contrast, it was restrained in tau expressing HEK293 cells. Interestingly, upon challenged with high glucose for 24 h, the levels of ROS in both of the cell lines all went back to normal level (**Fig. 5i**). These outcomes may because of the transcription of antioxygenic proteins, such as heme oxygenase-1 (HO-1) as reported by previous studies ^25,26^. Due to the proteome showed that the succinic dehydrogenase (SDH) A/B, which were less abundant in association with membrane in the hippocampus from hTau mice than that from Tau KO mice under hyperglycemia conditions (**Extended Data Fig. 8**). Thus, we further detected the activity of SDH, which is intimately related with not only electron transport chain but also TCA cycle ^27^. Surprisingly, the activity of SDH in tau expressing HEK293 cells was significantly greater than that in normal HEK293 cell, which may interpret the higher coupling efficiency in the former cells as observed in OCR. On the contrary, 1 h after meeting with high glucose, the activity of SDH in human tau expressing HEK293 cells was dramatically downregulated, and it was recovered 24 h after exposure to high glucose (**Fig. 5j**). These observations, therefore, demonstrated that human tau restricted the level of oxidative phosphorylation and it was also involved in regulating ROS production through mitochondrial respiration by influencing the activity of SDH.

## Conclusion

It has been well established that human tau is involved in regulating glucose metabolism and mitochondrial functions. However, as a microtubule binding protein, how does it contribute to these biological processes is not fully understood so far. Here, we showed that human tau promoted Warburg-effect like glycolytic metabolism under acute hyperglycemia conditions through modulating the homeostasis of protein-membrane association. Those proteins that were positively influenced by human tau for membrane association, including intermediate filament Kirts, heat shock proteins HSP90 and TRAP1, mitochondrial translational factor TFAM, and master signal transducer mTOR, showed close relations with glycolysis and mitochondrial homeostasis ^28,29^. Those were negatively influenced by human tau comprised electro transport chain proteins, ribosome proteins Rpls, as well as proteasome proteins Psmds (**Extended Data** Fig.9), indicating that except for influencing on oxidative phosphorylation, tau also participated in regulating protein synthesis and degradation. Of note, the dysfunction of ribosome and proteasome are also early events in Alzheimer’s Disease ^30–32^. These results are consistent with tau interactome studies ^33,34^, as well as the findings from analysis of tau evolution which indicated that human tau is involved in regulating protein-membrane association ^35^.

Human brain consumed up to 60% of all blood glucose ^36^. The processes of glycolysis and fatty oxidation require the rapid replenish of NAD^+^. Meanwhile, ROS generated as a by-product of these metabolisms need to be severely restrained. In the current study, we observed that under acute hyperglycemia conditions, tau not only promoted Akt activation but also facilitated aerobic glycolysis, which further recovered the equilibrium of NAD^+^ level. On the other hand, it restricted the level of oxidative phosphorylation, and reduced ROS production through modulating protein-membrane association (**summarized in Fig. 6**). These are according with the glycolytic advantage in cancer cell attributing to Warburg-effect ^37^. Thus, tau may counteract with the neurotoxicity induced by Aβ overproduction ^38^, such as overload of ROS and subsequent inflammation derived from microglia activation ^39^. However, the aggregation of tau may result in its disfunction on regulating protein-membrane associated, and subsequent energy metabolism deficit as seen in AD ^40^.

**Fig. 6.**
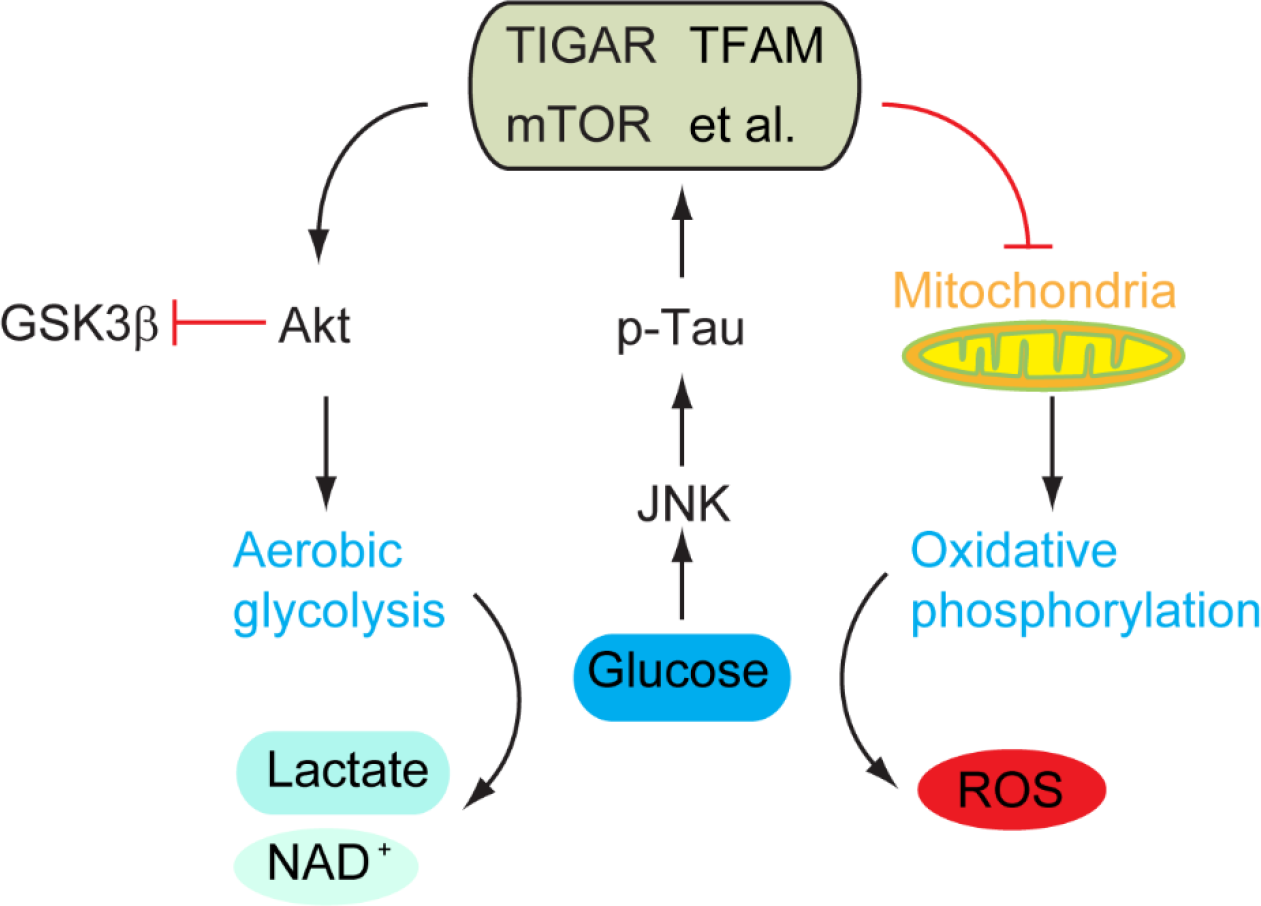
Schematic summary. High glucose/Acute glycemia conditions induced stress (i.e. ROS) triggered JNK activation, which further induced tau phosphorylation at multiple residues. Tau phosphorylation maintained the organization of membrane proteins, such as the membrane localization of TIGAR, TFAM, mTOR, TRAP1, electron transport chain, et al., Which subsequently strengthened aerobic glycolysis and restrained oxidative phosphorylation resembling with so called Warburg-effect.

The reprogramming of glycolytic metabolism in AD have been revealed in recent years. For example, the induced neuron from AD patient-derived fibroblasts exhibited Warburg-like metabolic transformation through elevating PKM2 expression and nuclear translocation ^41^. It has also been demonstrated that the lactate dependent histone modification promoted metabolism reprograming in AD model, which could facilitate the activation of microglia ^42^. However, the driving forces of the metabolic reprogramming in the neurodegenerative diseases are not clear. Based on the observations in this current study, we speculated that the phosphorylation of tau by JNK under physiological conditions plays important role in maintaining energy metabolism, particularly when blood glucose is elevated. However, the prolonged hyperphosphorylation of tau induced by other kinases and the aggregation of tau may give rise to the dysregulation of energy metabolism as well as the imbalance of protein synthesis and degradation in related neurogenerative diseases.

## Methods

### Animal

Human tau transgenic mice (B6.Cg-Tg(MAPT) 8cPdav *Mapt^tm^*^1^*^(EGFP)klt^/*J, Strain: 005491) and murine tau knockout (B6.129S4(Cg)-*Mapt^tm^*^1^*^(EGFP)klt^/*J, Strain: 029219) male mice were purchased from The Jackson Laboratory (Bar Harbor, ME, USA). C57 BL/6J mice were purchased from The Charles River. 3x Tg AD mice carrying the human mutations of APPswe, PS1M126V and TauP301L (B6; 129-Psen1^tm1Mpm^ Tg (APPSwe, tauP301L)1Lfa/Mmjax. Strain:034830) were acquired from Jackson Laboratory. Wild-type (WT) mice (B6129SF2/J, Strain: 101045) were used as controls for 3x Tg AD mice. All mice were housed in typical cages and lived in a humidity and temperature-normalized room with natural light-dark cycle which provided free water and food.

For acute hyperglycemia induction, Streptozotocin was diluted with citrate buffer (pH 4.2-4.5) to the final concentration (1%) before use. Four-month-old male mice of C57, Tau KO and hTau mice were received intraperitoneal injection (150mg/kg) and sacrificed 7 days after injection. Sex/age matched mice of each genotype injected with the same volume of PBS served as control. Similarly, 3x Tg-AD and WT mice were received same injection and sacrificed after 2 or 6 weeks.

### Chemicals and reagents

Streptozotocin (STZ) were acquired from Sigma-Aldrich (St. Louis, MO, USA). Fetal Bovine Serum (FBS), DMEM high-glucose medium and antibiotics (penicillin and streptomycin) were obtained from Gibco (Grand Island, NY, USA). Reactive oxygen species (ROS) test kit was provided by Nanjing Jiancheng Bioengineering Institute (Nanjing, China). Pierce BCA protein assay kit was provided by Thermo Fisher Scientific (Waltham, MA, USA). GIP, GLP-1 test kit were purchased from Jianglaibio (Shanghai, China). Pyruvate, Lactate and SDH assay kit were purchased from Elabscience (Wuhan, Hubei, China) .SP600125 (JNK inhibitor), INK-128 (mTORC1/2 inhibitor), Rapamycin (mTORC1 inhibitor) and JR-AB2-011 (mTORC2 inhibitor) were procured from MedChemExpress (Mercer County, NJ, USA)

### Cell culture

The HEK293 cell line that was stably transfected with the longest human tau (441 a. a.) cDNA ^43^ was provided by professor Jianzhi Wang from Tongji Medical College, Huazhong University of Science and Technology. Cells were cultured in DMEM supplemented with 10% FBS and penicillin/streptomycin (Gibco) and maintained in a humidified atmosphere with 5% CO2 at 37 ℃. Upon reaching 80% confluence, the cells were harvested and used for subsequent passage or high glucose with or without individual inhibitor treatment.

### Tissue/cell preparation and protein extraction

The mice were deeply anesthetized with isoflurane and euthanized. Brains were rapidly harvested, washed with normal saline and then separated into cortex and hippocampus. The cortex from right hemispheres in each group were homogenized by sonification for 20 seconds in radio immunoprecipitation assay (RIPA) lysis solution containing 1 mM phenylmethylsulphonyl fluoride (PMSF; Thermo Fisher Scientific, Waltham, MA, USA) supplemented with phosphatase and proteinase inhibitors. Cells were washed twice with PBS and homogenized by sonification in RIPA premixed with a cocktail of inhibitors as above. Subsequently, the homogenates were centrifuged at 12,000 rpm for 30 min at 4 ℃. The supernatants were collected and used for further experiments.

### Immunoblot analysis

Protein concentrations were determined by BCA protein assay kit. Tissue samples of equal amounts of proteins (20 µg) were loaded and separated on 8%, 10% and 12% SDS-PAGE gel and then transferred to polyvinylidene fluoride (PVDF) membranes (Millipore, Kankakee, IL, USA) in the transfer buffer. After transferring, the PVDF membranes were blocked with Blocking Buffer in tris-buffered saline containing 0.1% Tween 20 (TBST) for 30 min and then incubated with matched primary antibodies in TBST at 4 ℃. After binding with the corresponding primary antibody, the membranes were washed with TBST for 30 min three times and incubated with the secondary antibody for another 2 h at room temperature. After three washes in TBST for 30 min, the protein bands were visualized using chemiluminescence with an ECL kit (Advansta, Menlo Park, CA, USA), and the band intensities were analyzed using Image J software (NIH, Bethesda, MD, USA). β-Actin, β-tubulin and GAPDH was detected as an internal loading control.

### Tissue membrane protein extraction

Membrane and plasma extraction of Brain tissues were obtained with the Mem-PER Plus kit (Thermo fisher) according to the manufacturer’s protocols. The hippocampi from 4 month old male hTau (n=4), Tau KO (n=4), and C57 (n=4) that were treated with STZ for 7 dyas, and equal number controls of each group were used for membrane protein extraction in this study. The final extractions were collected and used for further experiments.

### TMT-labeled quantitative proteomic

Frozen hippocampi were quickly ground into fine and uniform powder in liquid nitrogen and then homogenized in 1 mL phenol extraction buffer, after that 1 mL saturated phenol with Tris-HCl (pH 7.5) was added. After several times shake, the mixture was kept at 4°C for 30 min. The upper phenolic phase was separated from the aqueous phase by centrifugation at 7100 g for 10 min at 4°C, transferred to a fresh tube and mixed with five volumes of pre-cold 0.1 M ammonium acetate-methanol. After being kept at −20 °C overnight, the mixture was centrifuged at 12,000 g for 10 min at 4°C to pellet precipitated protein. For the wash step, the pellet was resuspended twice with pre-cold methanol and twice with ice-cold acetone. Following another round of centrifugation, the pellet was collected, air-dried and resuspended with 300 μL lysate solution. After incubation of 3 h at room temperature, the solution was centrifuged to remove any insoluble fraction and the resulting supernatant contained the total extractable protein. The total protein concentrations were quantified by bicinchoninic acid assay.

According to the measured protein concentration, take the same quantity protein from each sample, and dilute different groups of samples to the same concentration and volume. Add 25mM DTT of the corresponding volume into the above protein solution to make the DTT final concentration about 5mM, and incubate at 55°C for 30-60 min. Then add the corresponding volume of iodoacetamide so that the final concentration was about 10mM, and place it in the dark for 15-30min at room temperature. Then 6 times of the volume of precooled acetone in the above system to precipitate the protein, and place it at − 20 °C for more than four hours or overnight. After precipitation, take out the sample and centrifuge at 8000 g for 10 min at 4 °C for collecting the precipitate. According to the amount of protein, add the corresponding volume of enzymolysis diluent (protein: enzyme = 50:1 (m/m), 100 μg of protein add 2 μg of enzyme) to redissolve the protein precipitate, then the solutions were incubated for digestion at 37°C for 12 h. Finally, samples were lyophilized or evaporated after enzymolysis.

For TMT/iTRAQ labelling, the lyophilized samples were resuspended in 30 μL 100mM TEAB and Labeling reaction in a 1.5 mL Ep tube. 20 μL acetonitrile was added to TMT/iTRAQ reagent vial at room temperature. The centrifuged reagents were dissolved for 5 min and mixed for centrifugation and repeat this step once. Then 10 μL of the TMT/iTRAQ label reagent was added to each sample for mixing. The tubes were incubated at room temperature for 1 h. Finally, 5 µL of 5% hydroxylamine were added to each sample and incubated for 15 min to terminate reaction. The labeling peptides solutions were lyophilized and stored at −80°C.

The Proteomic data analysis was performed by Shanghai Luming biological technology co., LTD (Shanghai, China). RP separation was performed on an 1100 HPLC System (Agilent) using an Agilent Zorbax Extend RP column (5 μm, 150 mm × 2.1 mm). Mobile phases A (2% acetonitrile in HPLC water) and B (90% acetonitrile in HPLC water) were used for RP gradient. The solvent gradient was set as follows: 0∼8 min, 98% A; 8∼8.01 min, 98%∼95% A; 8.01∼48 min, 95%∼75% A; 48∼60 min, 75∼60% A; 60∼60.01 min, 60∼10% A; 60.01∼70 min, 10% A; 70∼70.01 min, 10∼98% A; 70.01∼75 min, 98% A. Tryptic peptides were separated at an fluent flow rate of 300 μL/min and monitored at 210 nm. Samples were collected for 8-60 minutes, and eluent was collected in centrifugal tube 1-15 every minute in turn. Samples were recycled in this order until the end of gradient. The separated peptides were lyophilized for mass spectrometry.

All analyses were performed by a Q Exactive HF mass spectrometer (Thermo) equipped with a Nanospray Flex source (Thermo). Samples were loaded and separated by a C18 column (50 cm × 75 µm) on an EASY-nLCTM 1200 system (Thermo). The flow rate was 300 nL/min and linear gradient was 75 min (0∼50 min,2-28% B; 50∼60 min,28-42% B; 60∼65 min,42∼90%B;65∼75 min,90% B. mobile phase A = 0.1% FA in water and B = 0.1% FA in ACN). Full MS scans were acquired in the mass range of 350 – 1500 m/z with a mass resolution of 60000 and the AGC target value was set at 3e6. All MS/MS pattern acquisitions were fragmented with higher-energy collisional dissociation (HCD) with collision energy of 32. MS/MS spectra were obtained with a resolution of 45000 with an AGC target of 2e5 and a max injection time of 80 ms. The Q Exactive HF dynamic exclusion was set for 30.0 s and run under positive mode.

ProteomeDiscoverer (v.2.4.1.5) was used to search all of the raw data thoroughly against the Uniprot-Mus musculus-10090-2023.2.1. fasta database. Database search was performed with Trypsin digestion specificity. Alkylation on cysteine was considered as fixed modifications in the database searching. For protein quantification method, TMT/iTRAQ was selected. A global false discovery rate (FDR) was set to 0.01 and protein groups considered for quantification required at least 1 peptide.

Annotation of all identified proteins was performed using GO (http://www.blast2go.com/b2ghome; http://geneontology.org/) and KEGG pathway (http://www.genome.jp/kegg/). DEPs were further used for GO and KEGG enrichment analysis. Protein-protein interaction analysis was performed using the String (https://string-db.org).

### Determination of pyruvate, L-lactic acid levels

Brain tissues or cells were homogenized and centrifuged at 12,000 rpm for 20 min at 4 ℃. Then, the supernatant was collected and the concentration of total protein was determined with a BCA assay kit. The levels of pyruvate and L-lactic acid were evaluated with pyruvate and L-lactic acid assay kits according to the manufacturer’s protocols, respectively.

### Determination of NAD^+^ level

The levels of NAD+ in cells and tissues following glycolysis consumption were detected with a WST-8 kit (Beyotime, Shanghai, China). Briefly, cells or tissues were homogenized with NAD+/NADH extract from kit. Subsequently, the homogenates were centrifuged at 12,000 × g for 10 min at 4 ◦C. The supernatants were collected and split into two parts. One part of samples was heated to 60 ℃ to remove NAD^+^. Both parts and standards were incubated in ethanol dehydrogenase working solution in the dark for 20 min at 37 ℃. Then the samples were incubated in electronically coupled reagents in the dark for 10 min at 37 ℃. After incubation, the mean fluorescence intensity was detected at 450-nm using a spectra microplate reader (Thermo Fisher Scientific, Waltham, MA, USA).

### Oxygen consumption rate (OCR) and extracellular acidification rate (ECAR) analysis

OCR and ECAR were detected using seahorse XF24 analyzer (Seahorse bioscience, Billerica, MA, USA). Briefly, seeding HEK293 and HEK293-tau cells (2×10^4^ per well) in XF24 Cell Culture Microplates. Cells were covered with 475 μL assay medium (XF DMEM Medium, pH=7.4), 10 mM glucose, 1 mM sodium pyruvate and 1 mM L-glutamine. For OCR measurement, each port injection was processed with 1 μM oligomycin, 0.5 μM FCCP, 0.5 μM antimycin A and rotenone. For ECAR measurement, each port injection was performed with 50 μM rotenone and 0.5 μM 2-DG.

### Determination of ROS level

The levels of ROS in cells following OXPHOS process were assessed with a fluorescent probe 2′,7′-dichlorfluorescein-diacetate (DCFH-DA) kit. Briefly, the culture medium was removed and the cells were washed with PBS before the experiment. Then, the cells were cultured in serum-free medium containing 10 μM DCFH-DA dye in the dark for 20 min at 37 ℃. After incubation, cells were washed twice with PBS to eliminate the extracellular DCFH-DA. The fluorescence intensity was detected at 488-nm excitation and 535-nm emission wavelengths by an Accuri C6 flow cytometry (Beckman Coulter, Brea, CA, USA).

### SDH activity assay

Cells were homogenized with reagents from SDH activity assay kit and then centrifuged at 600 × g for 5 min at 4 ℃. Subsequently, the supernatants were collected and centrifuged at 15000 × g for 10 min at 4 ℃. Then the precipitates were sonicated for 5 min with new reagents from kit and centrifuged at 15000 g for 10 min. Finally, the supernatants were collected and the concentration of total protein was determined with a BCA assay kit. The SDH activity was detected at 600-nm by an Accuri C6 flow cytometry.

### Passivation of JNK and mTORC1/2 by inhibitor

The inhibitor was solubilized by DMSO (Yeasen, Shanghai, China) into an initial solution (the volume of DMSO was obtained by calculation from the MCE web site: https://www.medchemexpress.cn/molarity-calculator.htm). The supernatants of homogenates were collected after the cells were incubated for 24 h in high glucose medium (60 mM) containing the inhibitor. The working concentrations of each inhibitor were SP600125 (10 μM), INK-128 (12.5 μM), Rapamycin (20 μM) and JR-AB2-011 (1 μM)

### Statistical Analysis

All statistical analyses were conducted using GraphPad Prism 8.0 software (GraphPad, La Jolla, CA, USA). Results are presented as means ± s.e.m. Differences analyzed by utilizing two-tailed unpaired Student’s *t*-test or one-way or two-way ANOVA with Tukey’s post hoc test when appropriate. Significance was determined at *P*<0.05.

## Acknowledgement

We thank prof. Jianzhi Wang from Tongji Medical College, Huazhong University of Science and Technology and Dr. Jinwang Ye from College of Life Sciences and Oceanography, Shenzhen University for kindly providing human tau overexpressing HEK293 cell line.

## Data Availability Statement

The authors declare that the data supporting the findings of this study are available within the paper and the supplementary information files.

---

**Extended Data Fig. 1.**
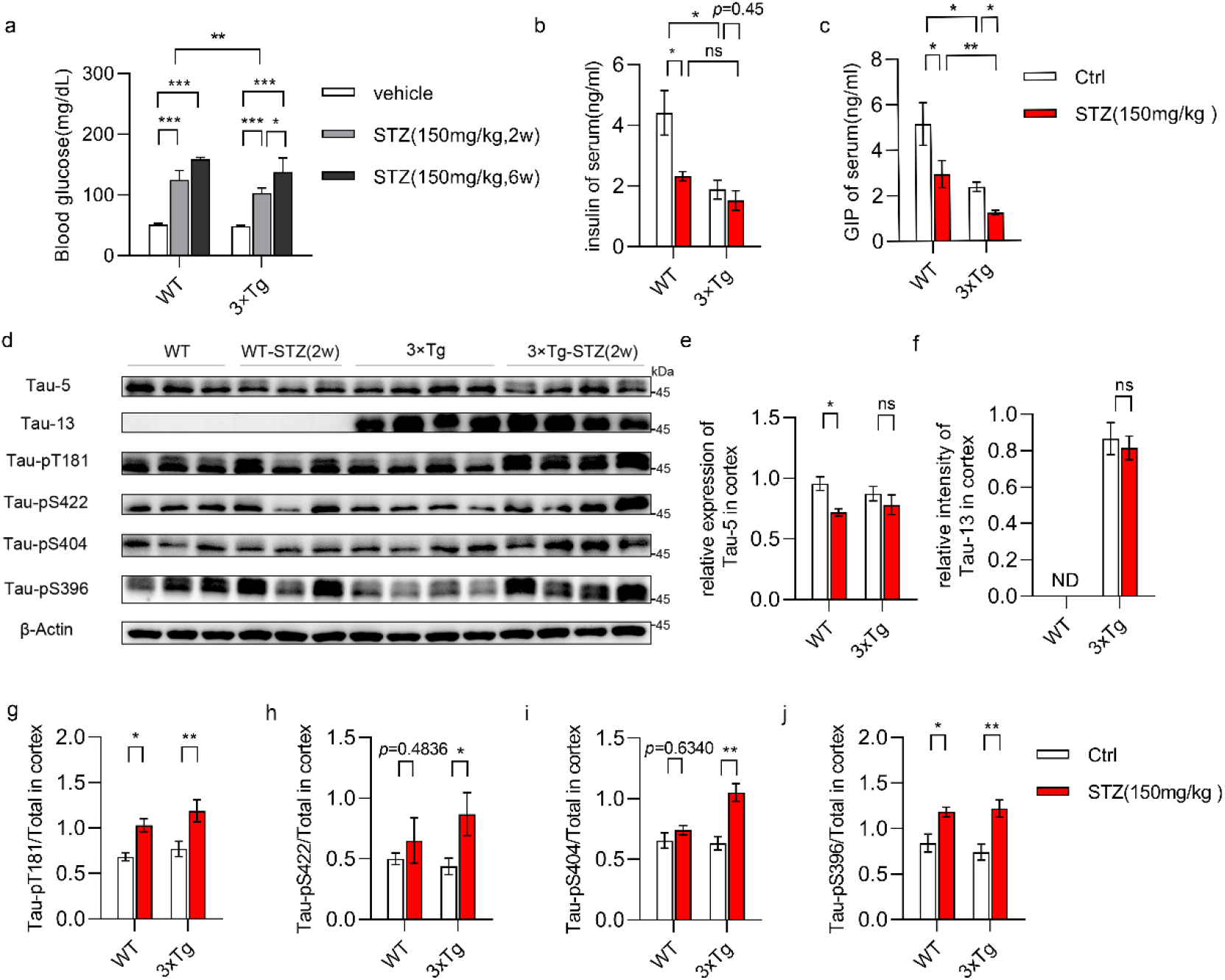
STZ injection induced tau phosphorylation in 3xTg AD mice. **a,** The fasting blood glucose levels of WT and 3xTg AD model mice at 4 months of age were measured (WT, n=14. 3x Tg AD, n=18), and then these mice were subjected to STZ treatment, thereafter, the fasting blood glucose were measured at 2 w (WT, n=8. 3x Tg AD, n=11) as well as at 6 w (WT, n=6. 3x Tg AD, n=7) after STZ injection, respectively. The levels of insulin (**b**) and GIP (**c**) in non-fasting serum of WT mice (n=3) and 3xTg AD model mice (n=6) that were injected with STZ (150 mg/kg) for 2 w, compared with that of WT mice (n=4) and 3xTg AD model mice (n=7) which were injected with PBS **d,** Immunoblot and quantifications of the protein levels of total tau (tau-5) (**e**), human tau (tau-13) (**f**), phosphorylated tau-pT181 (**g**), tau-pS422 (**h**), tau-pS404 (**i**), tau-pS396 (**j**) in the lysates of cortex from WT mice injected with PBS (n=3), or STZ (n=3) for 2 w, as well as 3xTg AD model mice injected with PBS (n=4), or STZ (n=4) for 2 w. \**P*<0.05, \*\**P*<0.01, \*\*\**P*<0.001 by two-way ANOVA with Tukey’s test (**a, b**) or two-tailed unpaired student’s *t*-test (**e-j**).

**Extended Data Fig. 2.**
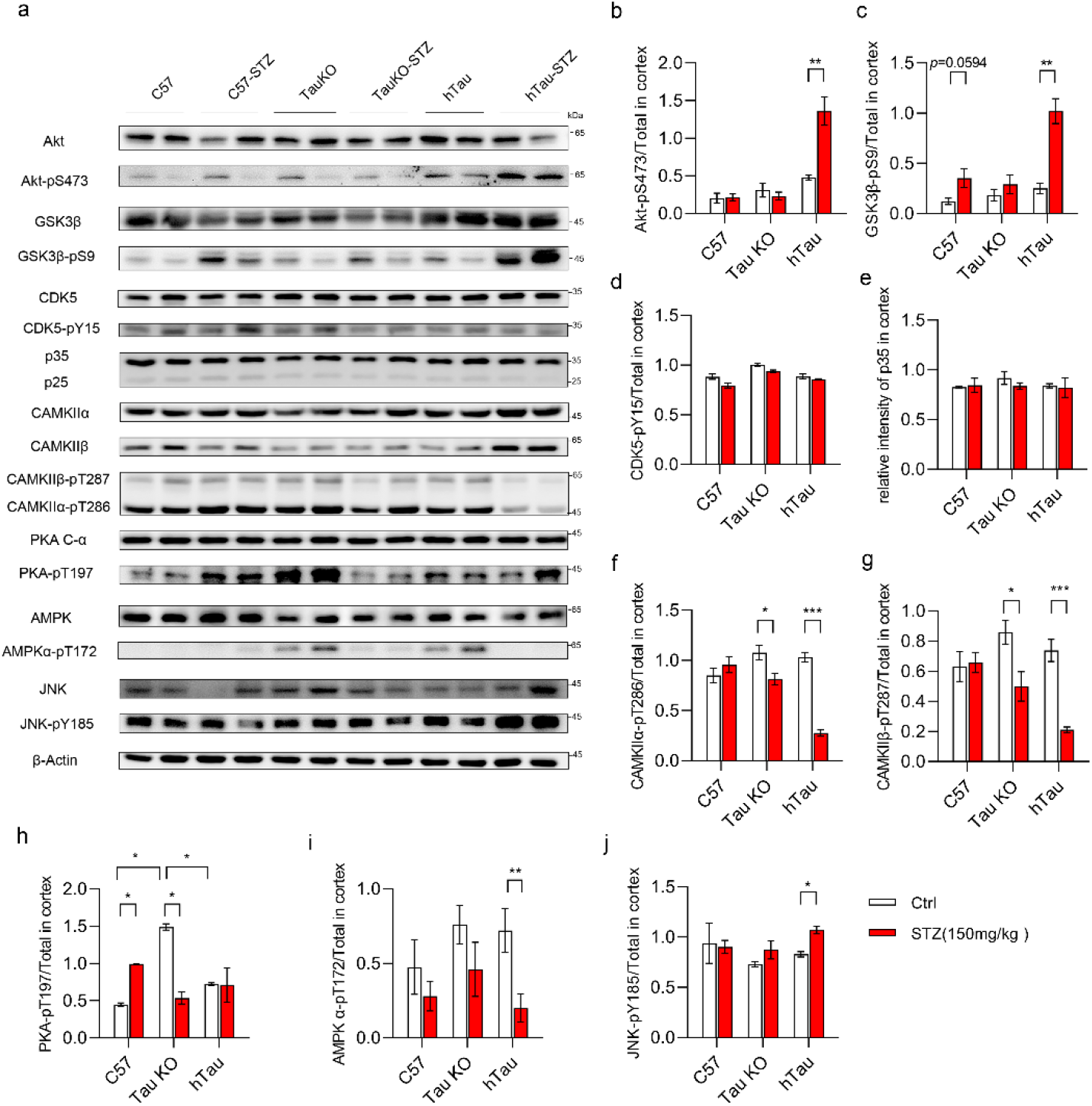
STZ injection induced JNK activation in hTau mice. **a,** Representative immunoblot of the cortex lysate 7 days after STZ injection (WT, n=4. Tau KO, n=4. hTau, n=4) compared with equal number controls. The ratio of Akt-pS437/Akt (**b**), GSK3β-pS9/tGSK3β (**c**), CDK5-pY15/CDK5 (**d)**, p35/actin (**e**), CAMKIIα-pT286/CAMKIIα (**f**), CAMKIIβ-pT287/CAMKIIβ (**g**), PKA-pT197/PKA (**h**), AMPK-pT172 /AMPK (**I**), and JNK-pY185/JNK (**j**) were analyzed and shown as the mean ±s.e.m., \**P*<0.05, \*\**P*<0.01, \*\*\**P*<0.001 by two-tailed unpaired student’s *t*-test (**b-g, i, j**) or two-way ANOVA with Tukey’s test (**h**).

**Extended Data Fig. 3.**
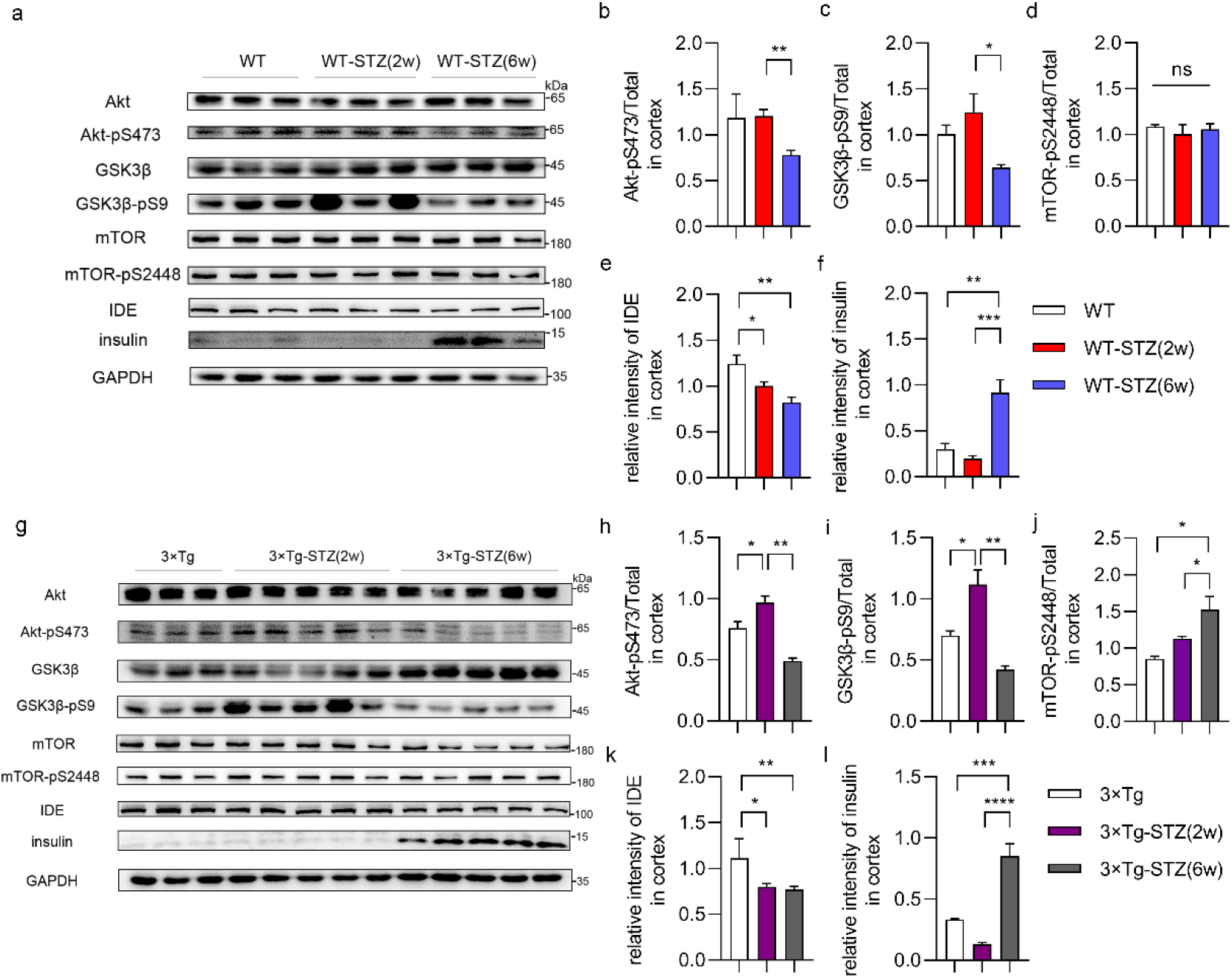
Prolonged hyperglycemia induced insulin resistance in brain. **a,** WT mice injected with STZ for 2 w (n=3), and 6 w (n=3), respectively, the lysates of cortex were isolated and subjected for **i**mmunoblot analysis compared with that of WT mice (n=3) injected with PBS for 2 w. Quantifications of the blot were shown as the mean ±s.e.m. of the ratios of Akt-pS473/Akt (**b**), GSK3β-pS9/GSK3β (**c**), mTOR-pS2448/mTOR (**d**), IDE/GAPDH (**e**), and insulin/GAPDH (**f**). **g**, 3x Tg AD mice were injected with STZ for 2 w (n=5), and 6 w (n=5), respectively, the lysates of cortex were isolated and subjected for **i**mmunoblot analysis compared with that of 3x Tg AD mice (n=3) injected with PBS for 2 w (n=3). Quantifications the blot were shown as the mean±s.e.m. of the ratios of Akt-pS473/Akt (**h**), GSK3β-pS9/GSK3β (**i**), and mTOR-pS2448/mTOR (**j**), IDE/GAPDH (**k**), and insulin/GAPDH (**l**), \**P*<0.05, \*\**P*<0.01, by one-way ANOVA with Tukey’s test.

**Extended Data Fig. 4.**
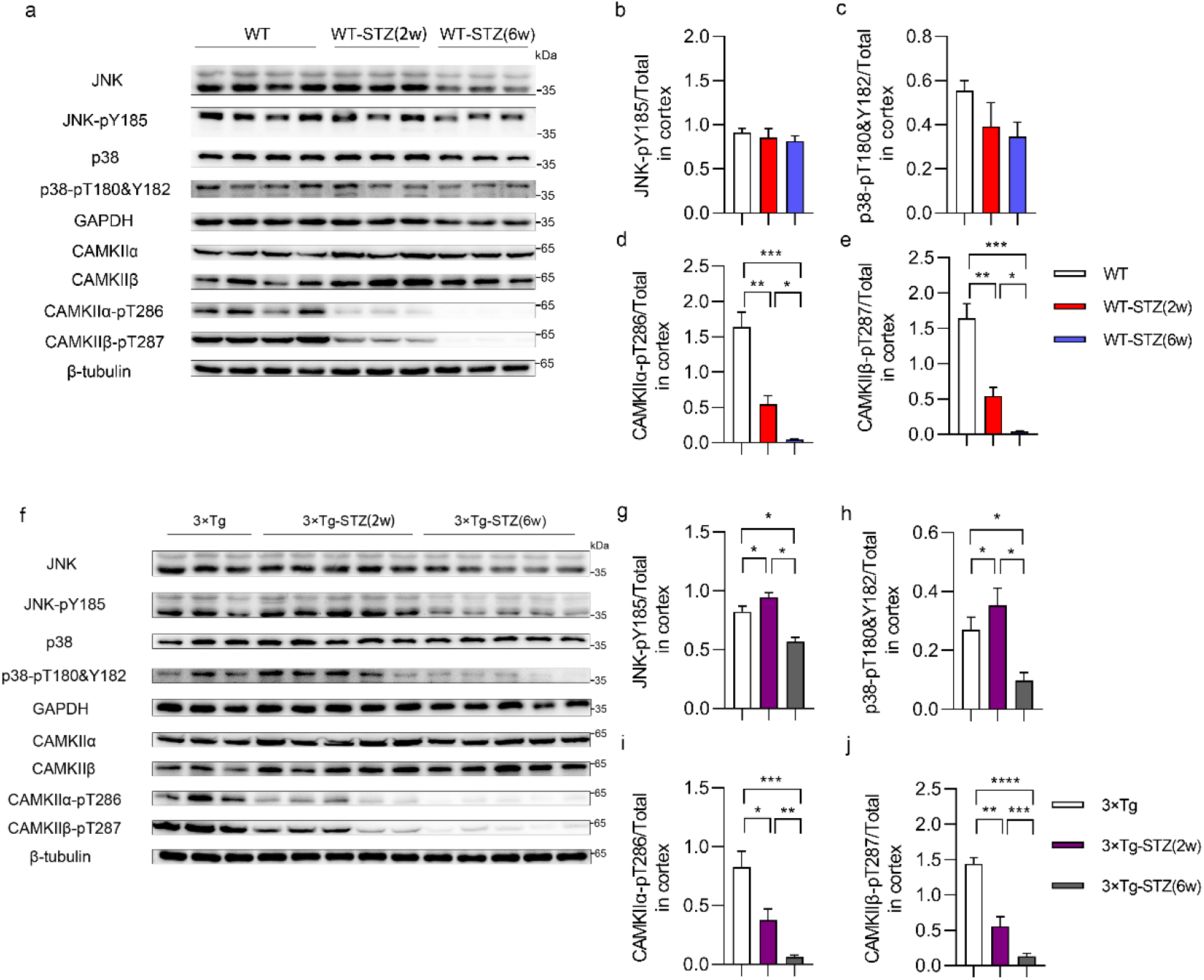
Prolonged hyperglycemia inhibited the phosphorylation of MAPKs and CaMKII. **a,** WT mice injected with STZ for 2 w (n=3), or 6 w (n=3), respectively, the lysates of cortex were isolated and subjected for **i**mmunoblot analysis compared with that of WT mice (n=4) injected with PBS for 2 w. Quantifications of the blot were shown as the mean ± s.e.m. of the ratios of JNK-pY185/JNK (**b**), p38-pT180&182/p38 (**c**), CAMKIIα-pT286/CAMKIIα (**d**), and CAMKIIβ-pT287/CAMKIIβ (**e**). **f**, 3x Tg AD mice were injected with STZ for 2 w (n=5), or 6 w (n=5), respectively, the lysates of cortex were isolated and subjected for **i**mmunoblot analysis compared with that of 3x Tg AD mice (n=3) injected with PBS for 2 w (n=5). Quantifications the blot were shown as the mean±s.e.m. of the ratios of JNK-pY185/JNK (**g**), p38-pT180&182/p38 (**h**), CAMKIIα-pT286/CAMKIIα (**i**), and CAMKIIβ-pT287/CAMKIIβ (**j**). \**P*<0.05, \*\**P*<0.01, by one-way ANOVA with Tukey’s test.

**Extended Data Fig. 5.**
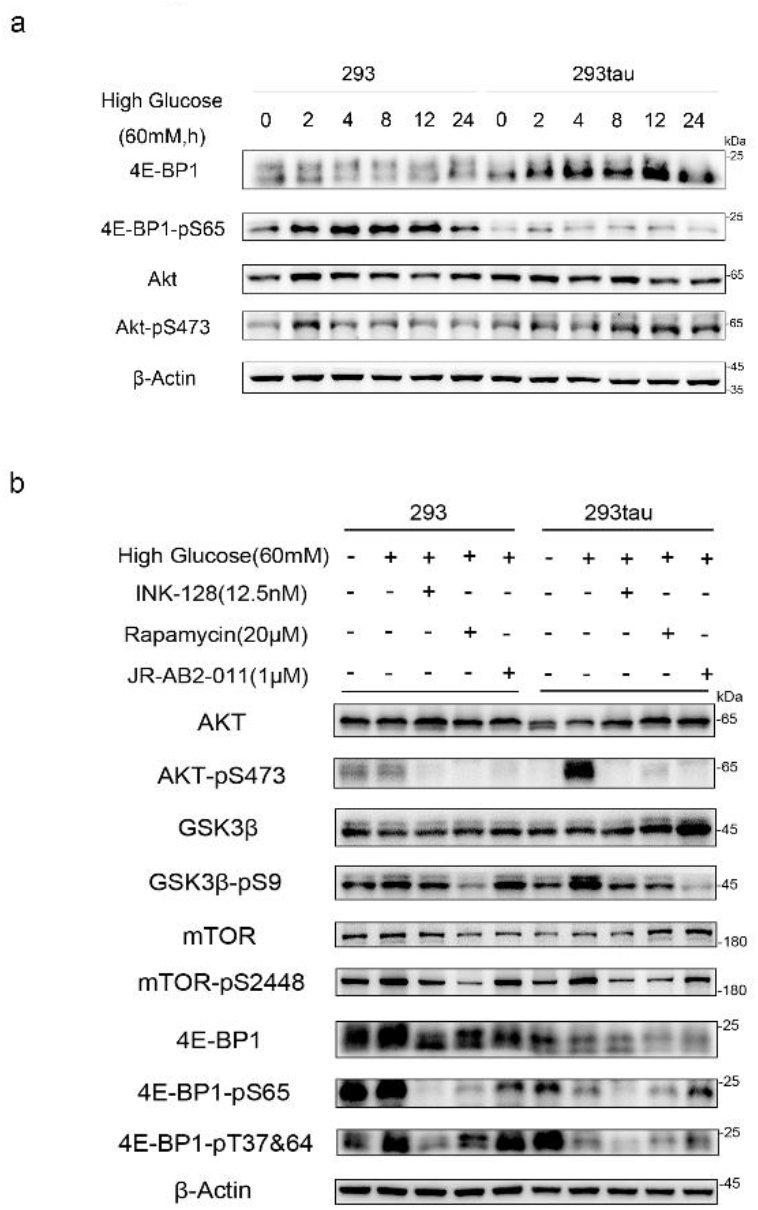
tau promoted alternative activation of mTORC2 under high glucose stress. **a**, Representative immunoblot of the normal HEK293 cells (293) and tau expressing HEK293 cells (293tau) that were cultured with high glucose 0-24 h. The protein level of 4E-BP1 and Akt, the phosphorylation level of 4E-BP1 at S65 and Akt at S473 were detected, actin served as a loading control. **b**, Representative immunoblot of the normal HEK293 cells (293) and tau expressing HEK293 cells (293tau) that were treated with high glucose (60 mM) for 24 h, with or without mTORC1/2 inhibitor INK-128 (12.5 nM), mTORC1 inhibitor rapamycin (20 μM), mTORC2 inhibitor JR-AB2-011 (1 μM). The protein level of Akt, GSK3β, mTOR, 4E-BP1, the phosphorylation level of Akt at S473, GSK3β at S9, mTOR at S2446, 4E-BP1 at S65 and at T37&64 were detected, actin served as a loading control.

**Extended Data Fig. 6.**
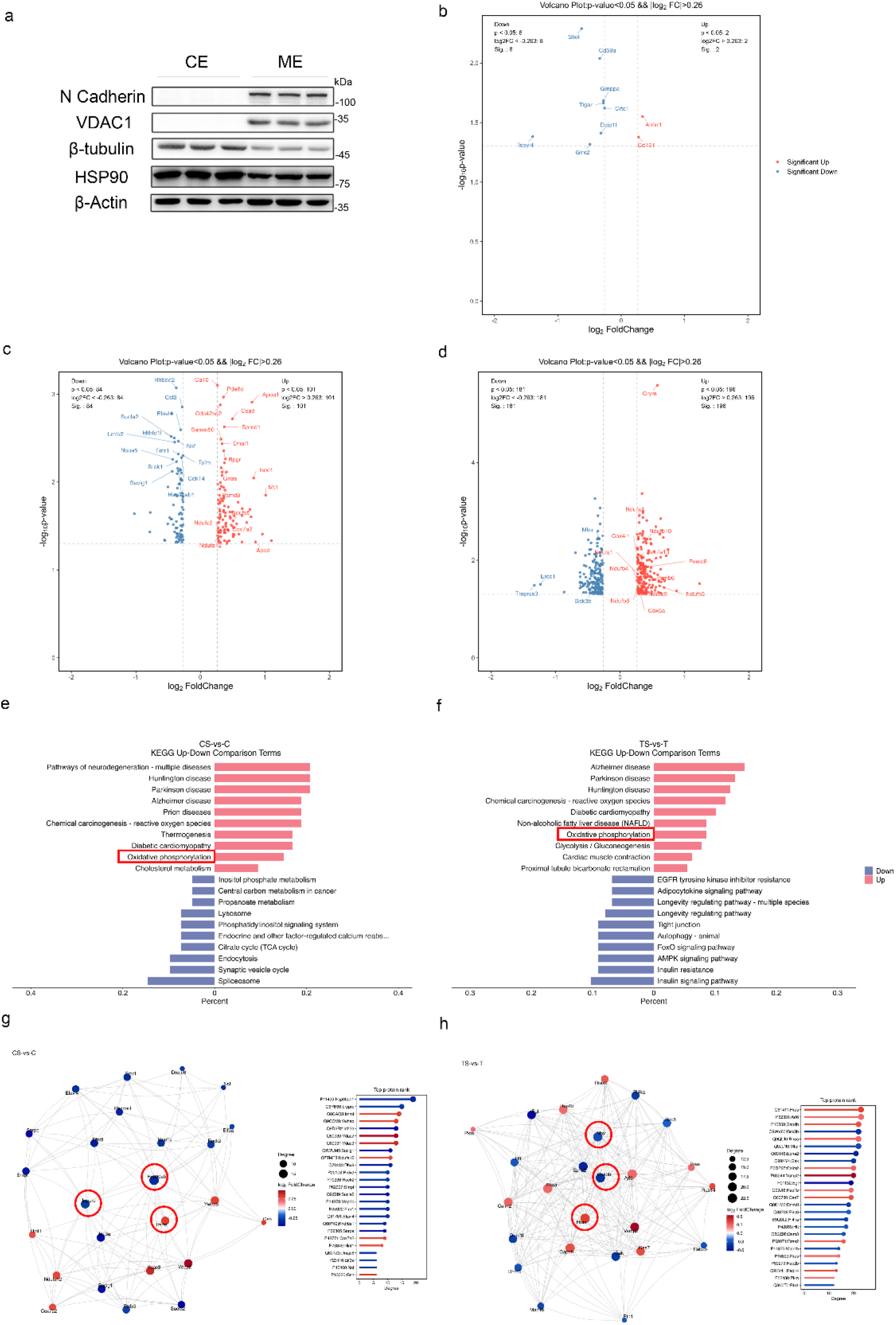
human tau maintained the homeostasis of protein membrane association in hippocampus under acute hyperglycemia conditions. **a**, Membrane associated proteins were isolated from the hippocampus and subjected to immunoblot for verification. The plasma membrane protein N-cadherin, and the mitochondrial membrane protein VDAC1 were only detected in the membrane extract, but not in the cytoplasmic extract. However, the cytoskeleton protein, β-tubulin and chaperon protein HSP90 were more enriched in cytoplasmic extract, indicating the membrane-associated proteins were isolated successfully. **b**, Volcano plots of the membrane-associated proteins (*P*<0.05, log_2_ Fold Change>0.26) in the hippocampus from 4 months old male mice (**b-d**). **b**, Acute hyperglycemia induced by STZ injection for 7 days elevated 2 proteins but depressed 8 proteins in association with membrane in hippocampus from hTau mice (n=4), respectively, comparing with not-treated groups (n=4). **c**, STZ treatment for 7 days increased 101 proteins, but reduced 84 proteins in association with membrane in the hippocampus of WT mice (n=4), respectively, comparing with non-treated WT mice (n=4). **d**, Under the same conditions, 196 proteins were upregulated, but 181 proteins were downregulated in the hippocampus of tau KO mice, respectively, comparing with non-treated tau KO mice. **e**, KEGG analysis showed that with respect to control WT mice, STZ-treatment facilitated neurodegenerative disease and oxidative phosphorylation, but restrained spliceosome related proteins in association with membrane. **f**, KEGG analysis showed that with respect to control tau KO mice, STZ treatment promoted neurodegenerative disease and oxidative phosphorylation, but weakened insulin signaling pathway related proteins in association with membrane. **g**, PPI network analysis revealed that HSP90ab1, LRPPRC, and IMMT are the top hub proteins in the network of discrepant membrane-associated between STZ treated WT mice and non-treated WT mice. **h**, PPI network analysis uncovered Hras, GSK3b, and mTOR are hub note of differential membrane-associated protein network between STZ treated tau KO mice and non-treated tau KO mice.

**Extended Data Fig. 7.**
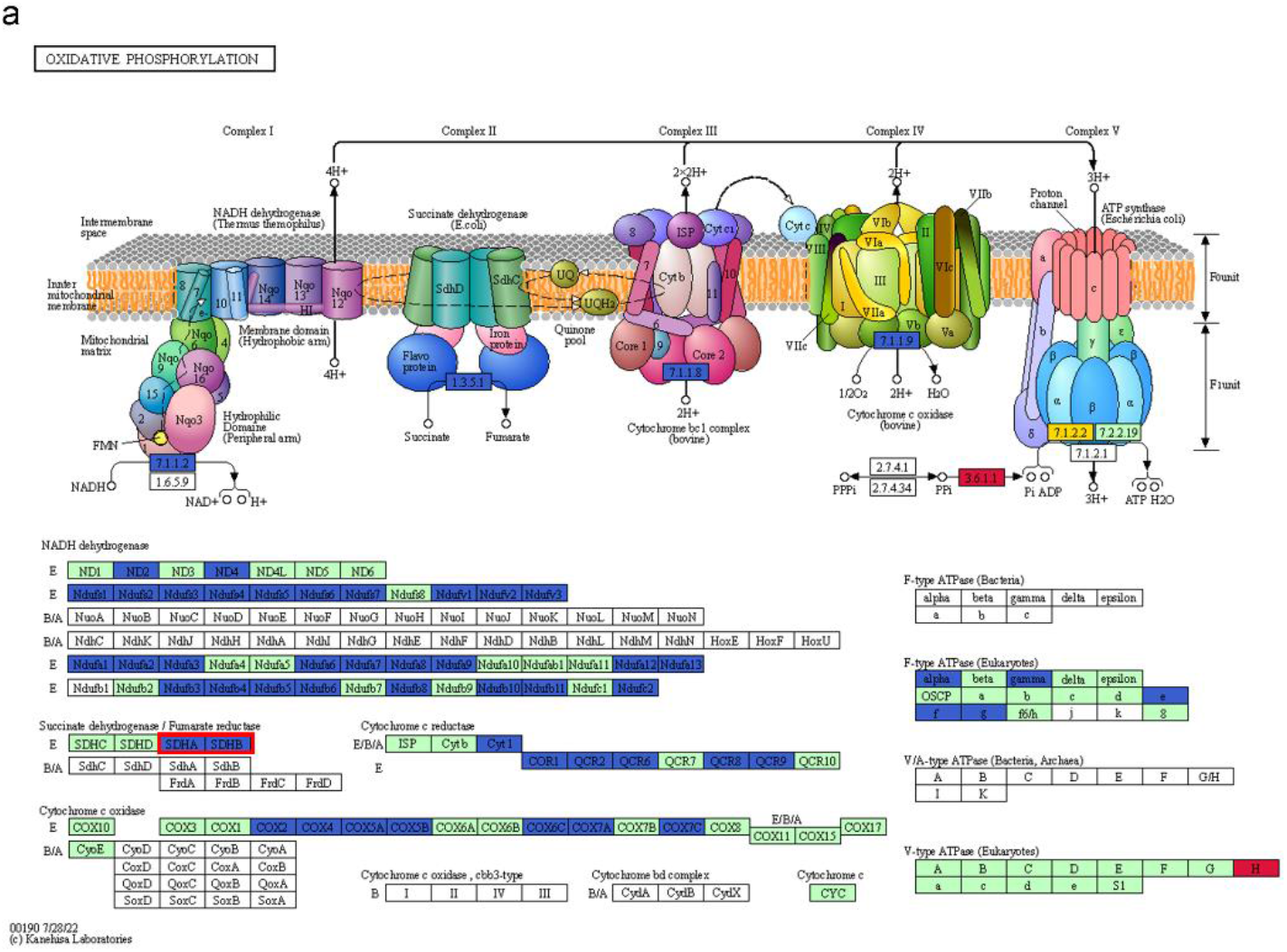
tau knockout resulted in dysregulation of the oxidative phosphorylation related protein locating on membrane under acute hyperglycemia conditions. KEGG analysis of membrane-associated proteome indicated that upon STZ treatment, more oxidative related proteins, including SDHA and SDHB were less abundant in association with membrane the hippocampus from hTau mice than that from Tau KO mice. The table with white background indicated the proteins that were not detected by iTRQ-MS. The table with green background indicated the proteins that did not show significant difference in the membrane extract of hippocampus between hTau mice and Tau KO mice. The table with blue background indicated the proteins that were significant less abundant in association with membrane in the hippocampus between hTau mice and Tau KO mice. The table with red background indicated the proteins that were significant more abundant in association with membrane in the hippocampus between hTau mice and Tau KO mice.

**Extended Data Fig. 8.**
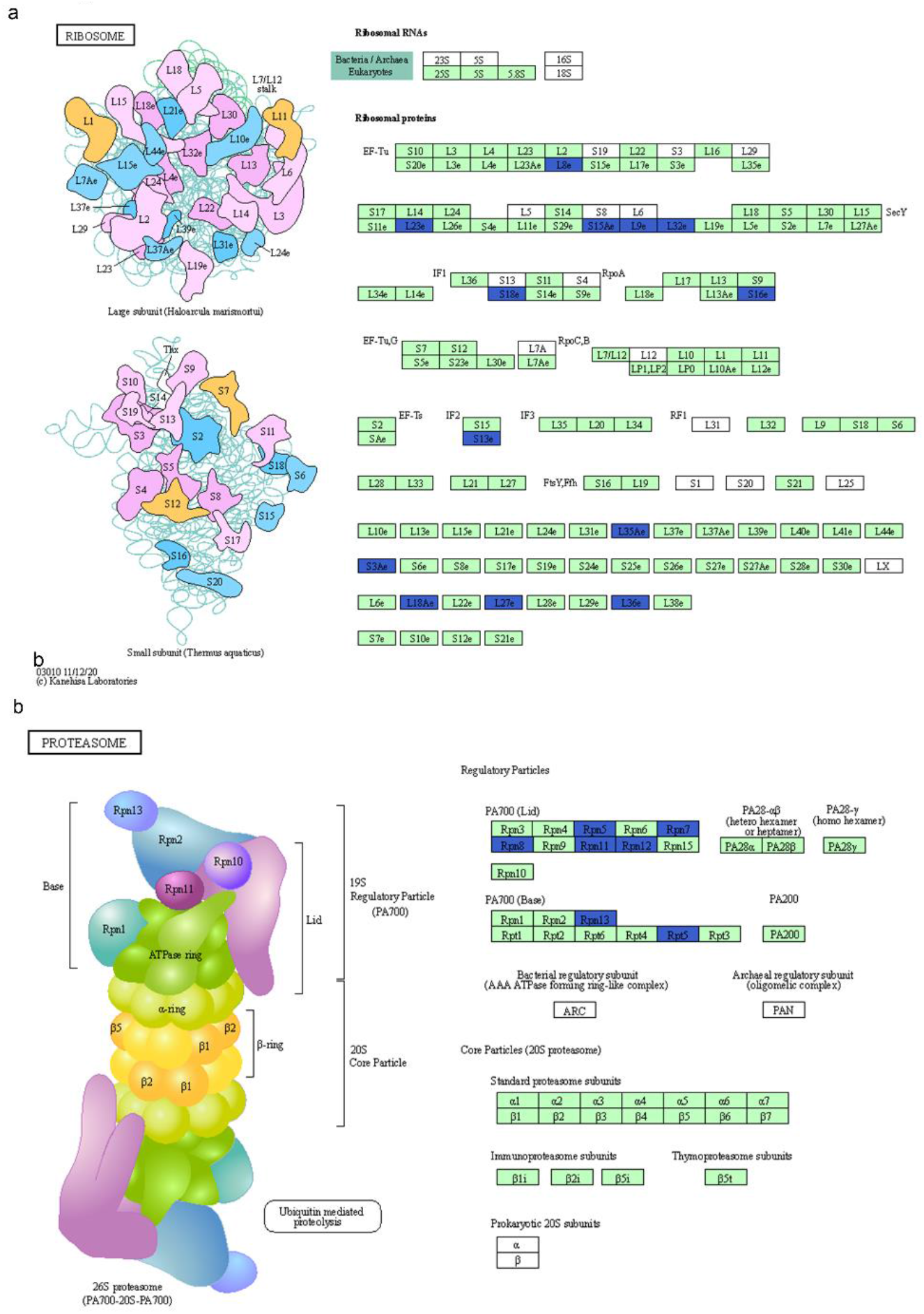
human tau arrested the association of the of 26s proteasome and ribosome related proteins with membrane. KEGG analysis of membrane-associated proteome indicated that the components of ribosome (**a**) and proteasome (**b**) were less abundant in association with membrane in the hippocampus from hTau mice than that from Tau KO mice. The table with green background indicated the proteins that did not show significant difference in the membrane extract of hippocampus between hTau mice and Tau KO mice. The table with blue background indicated the proteins that were significant less abundant in association with membrane in the hippocampus between hTau mice and Tau KO mice.

**Extended Data Fig. 9.**
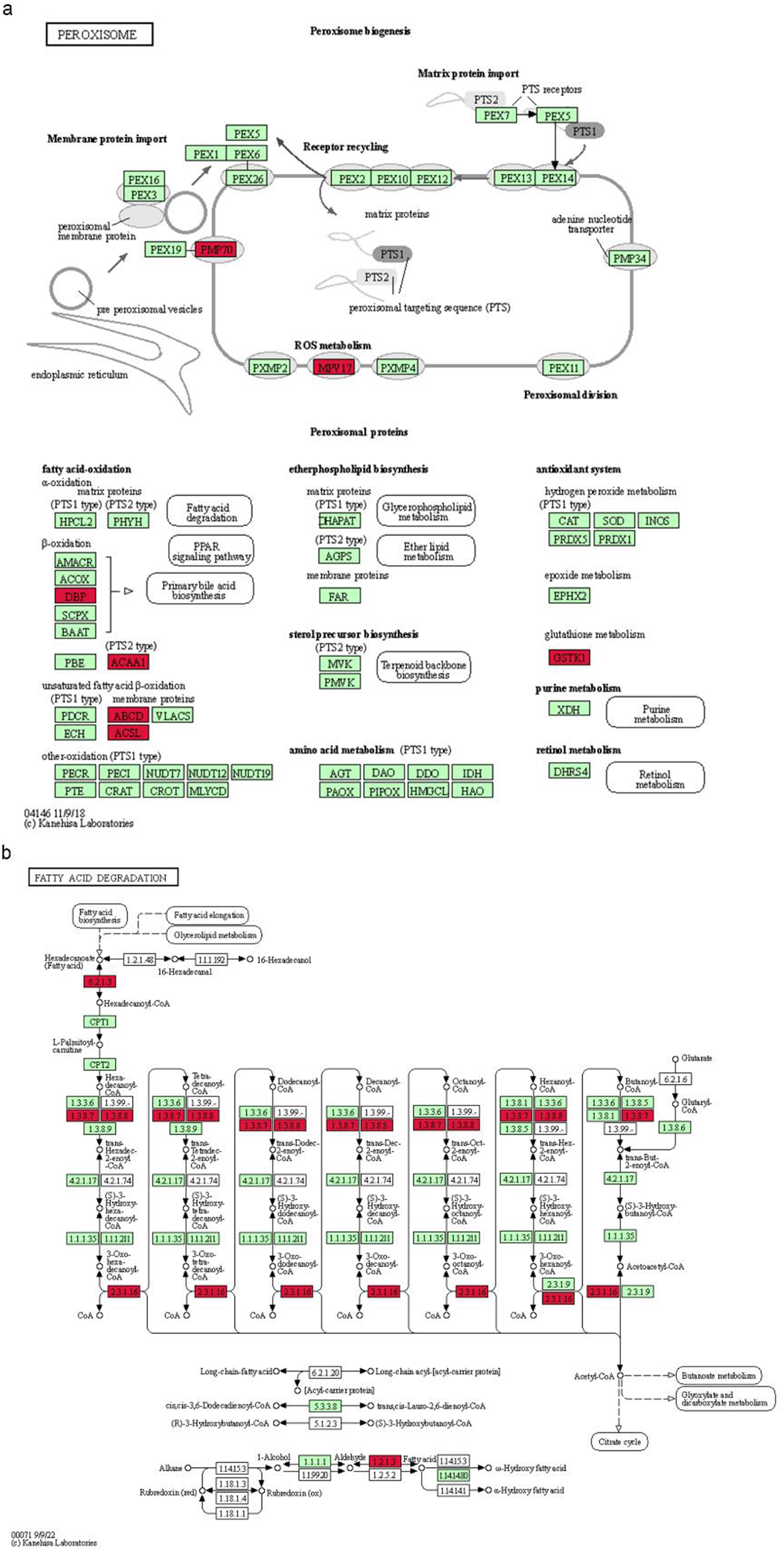
human tau augment the association of the components of peroxisome and fatty acid degradation enzyme. KEGG analysis membrane-associated proteome indicated that the components of peroxisome (**a**) and the enzymes that are involved in fatty acid degrading (**b**) were more abundant in association with membrane in the hippocampus from hTau mice than that from Tau KO mice. Significantly different proteins were indicated with red. The table with green background indicated the proteins that did not show significant difference in the membrane extract of hippocampus between hTau mice and Tau KO mice. The table with red background indicated the proteins that were significant more abundant in association with membrane in the hippocampus between hTau mice and Tau KO mice.6.2.1.3: acyl-CoA synthetase long-chain family member 1, 1.3.8.7: acyl-Coenzyme A dehydrogenase, 1.3.8.8: acyl-Coenzyme A dehydrogenase, 2.3.1.16: acetyl-Coenzyme A acyltransferase 1A, 1.2.1.3: aldehyde dehydrogenase family 7, member A1.

